# Cell-free Trim-Away reveals the mechanism of antibody-mediated protein degradation by TRIM21

**DOI:** 10.1101/2022.07.23.501259

**Authors:** Tycho E.T. Mevissen, Anisa V. Prasad, Johannes C. Walter

## Abstract

TRIM21 is a cytosolic antibody receptor and E3 ubiquitin ligase that promotes destruction of a broad range of pathogens. TRIM21 also underlies the antibody-dependent protein targeting method Trim-Away. Current evidence suggests that TRIM21 binding to antibodies leads to formation of a self-anchored K63 ubiquitin chain on the N-terminus of TRIM21 that triggers the destruction of TRIM21, antibody, and target protein. Here, we report that addition of antibody and TRIM21 to *Xenopus* egg extracts promotes efficient degradation of endogenous target proteins, establishing cell-free Trim-Away as a powerful tool to interrogate protein function. Chemical methylation of TRIM21 had no effect on target proteolysis, whereas deletion of all lysine residues in targets abolished their ubiquitination and proteasomal degradation. These results demonstrate that target protein but not TRIM21 polyubiquitination is required for Trim-Away, and they suggest that current models of TRIM21 function should be fundamentally revised.

## INTRODUCTION

Tripartite motif containing protein 21 (TRIM21) is an intracellular immunoglobulin receptor and E3 ubiquitin ligase that functions in the innate immune response (Kiss and James, 2021). When non-enveloped viruses enter a cell, they bring with them capsid-bound serum antibodies (Mallery et al., 2010). TRIM21 binds to these antibodies, leading to activation of its E3 ubiquitin ligase activity. TRIM21-dependent ubiquitination then promotes destruction of the virus via a mechanism that relies on the proteasome and the p97 ATPase (Hauler et al., 2012). During this process, the antibody and TRIM21 are also destroyed. The ubiquitination activity of TRIM21 also stimulates proinflammatory immune signaling pathways (McEwan et al., 2013). Thus, TRIM21 forms an important barrier against pathogens by bridging the adaptive and innate immune responses.

The domain architecture and enzymology of TRIM21 suggest that activation of its E3 ubiquitin ligase activity requires higher order assembly (Kiss and James, 2021). TRIM21 contains an N-terminal RING domain and an internal coiled-coil domain that mediates anti-parallel TRIM21 homodimerization, placing the RING domains at opposite ends of a TRIM21 dimer (Figure 1A). Because RING domain dimerization is often required for productive interaction with E2 ubiquitin-conjugating enzymes, the TRIM21 homodimer is believed to be inactive. TRIM21 also contains a C-terminal PRYSPRY domain that, upon viral entry, binds tightly to the Fc portion of antibodies coating the viral capsid (James et al., 2007) (Figure 1A). In the case of adenovirus, at least 24 TRIM21 homodimers must bind the viral particle to achieve full virus neutralization (Zeng et al., 2021). Collectively, these observations suggest that TRIM21 activation depends on juxtaposition of RING domains from adjacent TRIM21 homodimers bound to antibody-decorated pathogens. This clustering mechanism is attractive because it ensures that TRIM21 is only activated on multivalent, antibody-bound structures such as viruses.

**Figure 1.**
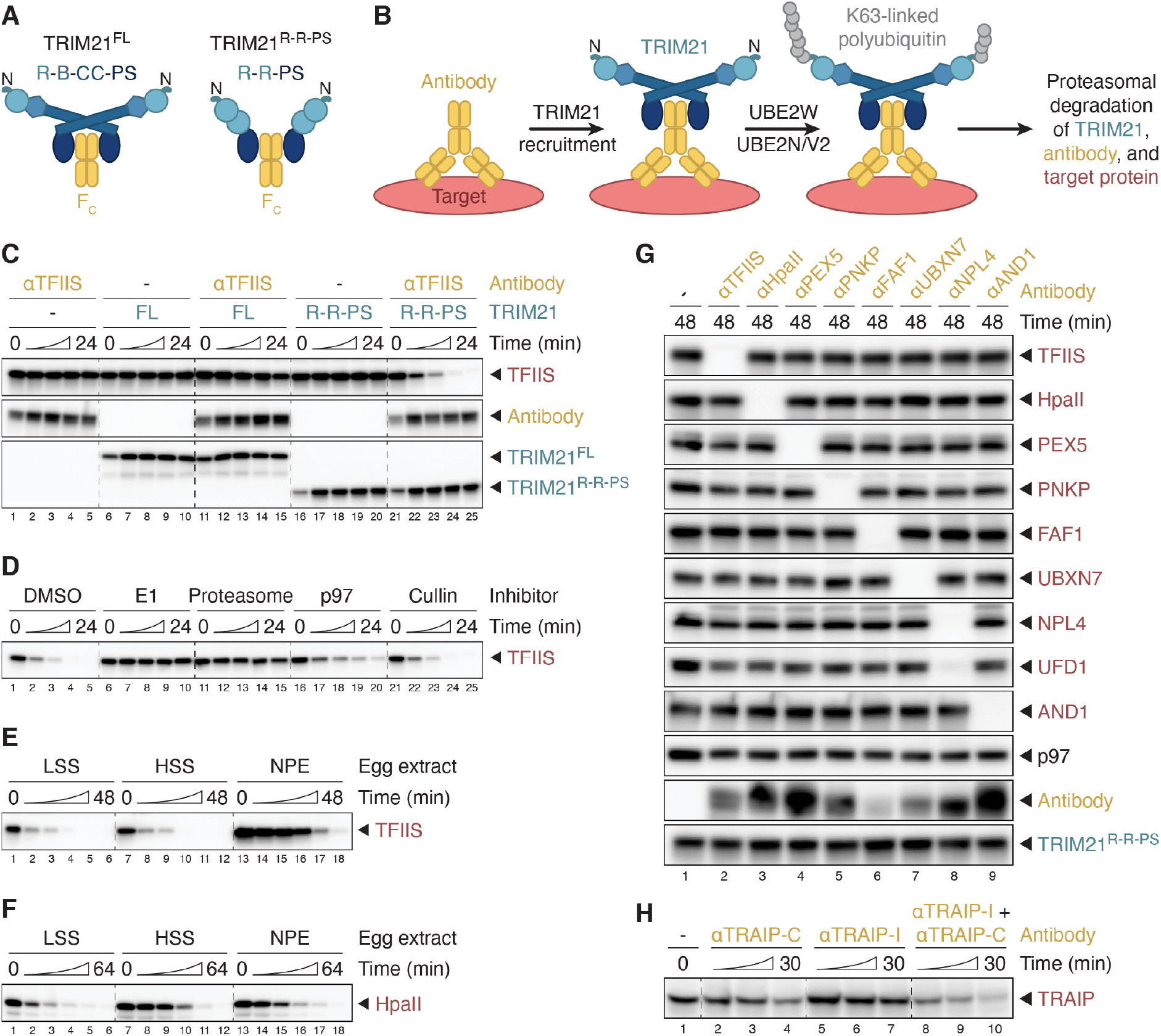
Cell-free Trim-Away in *Xenopus* egg extracts. **(A)** Domain architectures of full-length (FL) TRIM21 and a minimal TRIM21 construct (TRIM21^R-R-PS^). R, RING domain; B, B-box domain; CC, coiled-coil domain; PS, PRYSPRY domain; F_c_, antibody fragment. **(B)** Simplified schematic of the current Trim-Away model. The protein of interest (red) is bound by an antibody (yellow), which in turn is recognized by TRIM21 (shades of blue). Subsequent TRIM21 activation results in sequential mono- and polyubiquitination of its N-terminus with K63-linked polyubiquitin, which requires the E2 ubiquitin-conjugating enzymes UBE2W and UBE2N/UBE2V2, respectively. The polyubiquitin chain on TRIM21 has been proposed to promote the proteasomal degradation of all three components. **(C)** Trim-Away of endogenous TFIIS in *Xenopus* egg extract (HSS, high-speed supernatant) comparing recombinant human TRIM21^FL^ and TRIM21^R-R-PS^ (final concentrations of 1 µM). **(D)** Trim-Away of TFIIS in HSS using TRIM21^R-R-PS^ in the presence of inhibitors targeting E1 enzyme (MLN7243), the proteasome (MG-262), p97 (NMS-873), or the NEDD8 activating enzyme to inhibit Cullin-RING E3 ligases (MLN4924). **(E)** Endogenous TFIIS was targeted by TRIM21^R-R-PS^ in three types of *Xenopus* egg extract: LSS (low-speed supernatant), HSS (high-speed supernatant), and NPE (nucleoplasmic extract). Note that TFIIS is more abundant in NPE than in LSS and HSS. **(F)** Egg extracts were supplemented with 250 nM recombinant HpaII, a bacterial DNA methyltransferase, to compare Trim-Away efficiencies using TRIM21^R-R-PS^ in LSS, HSS, and NPE at a constant target protein concentration. **(G)** The specificity of cell-free Trim-Away was assessed by using a panel of polyclonal antibodies targeting various endogenous or exogenous (HpaII, supplemented to 250 nM) substrates in an end-point assay. All antibodies were raised against entire proteins or protein domains. UFD1 is a constitutive binding partner of the targeted protein NPL4. p97 was visualized because several tested Trim-Away substrates (i.e. FAF1, UBXN7, and NPL4) are p97 interactors. **(H)** Endogenous TRAIP was targeted by TRIM21^R-R-PS^ in NPE comparing two polyclonal peptide antibodies: one raised against the C-terminus (TRAIP-C), and one against an internal fragment (TRAIP-I). The samples of experiments in panels C-F were treated at indicated times with inhibitors targeting E1 and the proteasome as well as the non-specific deubiquitinase USP2 for 30 min prior to western blot analysis. Untreated samples of identical assays are shown in Figures S1B-E.

The current model of TRIM21 activation envisions that after clustering-induced RING-dimerization, TRIM21 interacts with two different E2 enzymes (Figure 1B) (Fletcher et al., 2015). UBE2W first promotes monoubiquitination of TRIM21’s N-terminus; UBE2N/UBE2V2 then builds a K63-linked polyubiquitin chain on the Nterminal monoubiquitin. However, for steric reasons, monoubiquitinated TRIM21 can only undergo UBE2N/UBE2V2-dependent polyubiquitination in trans. Therefore, TRIM21 ubiquitination requires the juxtaposition of at least three TRIM21 RING domains (Kiss et al., 2021). The K63-linked polyubiquitin chain on TRIM21 has been proposed to trigger immune signaling as well as the proteasomal degradation not only of TRIM21, but also of antibody and substrate, whose ubiquitination has not been detected and is assumed not to occur (Figure 1B) (Fletcher et al., 2015). The destruction of proteins without direct ubiquitination is unconventional, but would allow TRIM21 to destroy a potentially unlimited number of substrates, as long as they are bound by a critical number of antibodies (Kiss and James, 2021; Zeng et al., 2021). Because the proteasome generally does not act on K63-linked polyubiquitin chains, it has been proposed that this modification scaffolds subsequent formation of K48 chains, the canonical proteasomal degradation signal. Indeed, TRIM21 overexpression in cells leads to formation of TRIM21-linked heterotypic K48/K63 chains (Fletcher et al., 2015). Many aspects of this TRIM21 activation model are based on biochemical and structural experiments lacking antibody or substrate, and whether substrate ubiquitination is required for TRIM21 function has never been directly examined.

TRIM21 has been repurposed to allow antibody-mediated degradation of endogenous proteins, a method called Trim-Away (Clift et al., 2017, 2018). In Trim-Away, antibodies directed against a protein of interest are delivered into cells via micro-injection or electroporation. Endogenous (or delivered) TRIM21 then promotes ubiquitin-mediated proteolysis of the target, which is accompanied by antibody and TRIM21 destruction. Because Trim-Away works with commercially available antibodies, requires no genetic modification of the target, and leads to rapid protein depletion, Trim-Away is a powerful tool to modulate protein levels. As for TRIM21dependent virus neutralization, the relevant targets of ubiquitination are unknown. Although Trim-Away has been successfully applied in diverse cell types and embryos (Chen et al., 2019; Clift et al., 2017; Israel et al., 2019; Weir et al., 2021), it has not been used in cell-free extracts.

Cellular extracts are powerful systems to study complex biochemical processes that have not been reconstituted with purified components. *Xenopus laevis* egg extracts are especially versatile, having led to important advances in our understanding of many cellular processes, including DNA replication, DNA repair, mitosis, cell cycle progression, organelle assembly, apoptosis, and cytoskeletal dynamics (Hardwick and Philpott, 2015; Hoogenboom et al., 2017). To identify a protein’s function in a cell-free system, the protein is usually removed from the extract using immobilized antibodies (“immunodepletion”). However, this procedure has several drawbacks: it can cause extract inactivation due to dilution; it cannot remove proteins that are part of larger cellular structures, such as integral membrane proteins; it cannot be used to rapidly remove a protein at an intermediate step of a complex biochemical pathway, which is also a problem for investigating protein function by genetic strategies; finally, immunodepletion is time-consuming and requires large amounts of antibody. For these reasons, we sought to adapt Trim-Away for use in cell-free extracts.

Here, we show that addition of recombinant TRIM21 and antibodies to egg extracts is sufficient to induce rapid and efficient degradation of diverse, endogenous and exogenous proteins, and we demonstrate that depleted extract can be functionally reconstituted with recombinant protein. Furthermore, we use this system to explore the mechanism by which TRIM21 targets proteins for proteasomal destruction. We demonstrate that TRIM21dependent substrate proteolysis requires direct target protein ubiquitination on lysine residues, whereas TRIM21 and antibody ubiquitination are dispensable. Finally, we show that TRIM21 does not ubiquitinate its Nterminus during cell-free Trim-Away, and that it forms heterotypic K48- and K63-linked polyubiquitin chains on its substrates. Our results expand the toolkit for studying protein function in cell-free extracts, and they suggest that current models of TRIM21 action should be revised.

## RESULTS

### Cell-free Trim-Away in *Xenopus* egg extracts

To adapt Trim-Away for use in *Xenopus* egg extracts, we first attempted to target endogenous TFIIS (a transcription elongation factor) in a high-speed supernatant (HSS) of egg lysate using an affinity-purified polyclonal antibody. Addition of TFIIS antibody to extract did not promote TFIIS degradation, suggesting that endogenous TRIM21 levels were not sufficient to support Trim-Away (Figure 1C, lanes 1-5). Moreover, supplementing HSS with recombinant human full-length TRIM21 (referred to as TRIM21^FL^) (Figure S1A) together with TFIIS antibody promoted inefficient target proteolysis within a short time course of up to 24 minutes (Figure 1C, lanes 11-15). To circumvent the necessity of TRIM21 clustering to promote RING dimerization and activation, a minimal TRIM21 construct containing the PRYSPRY domain fused to two RING domains (TRIM21^R-R-PS^, Figures 1A and S1A) was developed previously (Kiss et al., 2021). Strikingly, TRIM21^R-R-PS^ was able to trigger fast and efficient antibody-dependent destruction of TFIIS in HSS (Figure 1C, lanes 21-25), establishing cell-free Trim-Away.

TRIM21-dependent target destruction in cells requires ubiquitin-mediated proteasomal degradation. Accordingly, the E1 inhibitor MLN7243 or the proteasome inhibitor MG-262 completely abrogated Trim-Away of TFIIS in egg extract (Figure 1D). Furthermore, p97 inhibition with NMS-873 delayed TFIIS proteolysis, suggesting that p97 may assist the proteasome in efficient substrate destruction. In contrast, a Cullin-RING E3 ligase inhibitor (MLN4924) did not impair Trim-Away, as expected (Figure 1D). Together, our data suggest that cell-free Trim-Away occurs by a mechanism analogous to that seen in human cells.

TRIM21-mediated TFIIS degradation was not limited to HSS, but also occurred in unfractionated egg lysate (LSS, low-speed supernatant) and a nucleoplasmic extract (NPE) derived from *Xenopus* eggs (Figure 1E). However, TFIIS was seemingly degraded with slower kinetics in NPE compared to HSS and LSS. Since TFIIS is also more abundant in NPE, we supplemented all three extracts with the same amount of an exogenous protein, the bacterial DNA methyltransferase HpaII. As shown in Figure 1F, in the presence of a HpaII antibody, Trim-Away of HpaII proceeded with more comparable kinetics in LSS, HSS, and NPE, demonstrating that cell-free Trim-Away is efficient in all tested egg extract types.

We next assessed the specificity of cell-free Trim-Away by targeting a wide range of proteins. TRIM21^R-R-PS^ alone did not promote degradation of any of the eight proteins of interest, whereas co-supplementation of polyclonal antibodies resulted in rapid, specific, and complete substrate degradation (Figure 1G). Interestingly, the constitutive binding partner UFD1 of the targeted protein NPL4 was also efficiently destroyed (Figure 1G, lane 8). The UFD1-NPL4 heterodimer is a cofactor of the p97 ATPase, yet neither p97 levels nor levels of two other cofactors (FAF1 or UBXN7) were affected by Trim-Away with NPL4 antibody. Thus, direct and stable interactors of targeted proteins may be co-depleted during Trim-Away.

All polyclonal antibodies used in Figure 1G were raised against entire proteins or protein domains, and we noticed that such antibodies often promoted rapid substrate degradation. In contrast, polyclonal antibodies recognizing short peptides, such as C-terminal or internal fragments of the E3 ubiquitin ligase TRAIP (Figure 1H, lanes 1-7), were generally less efficient or even completely inactive for cell-free Trim-Away, suggesting that multiple epitopes need to be available. Consistent with this interpretation, we observed more effective TRAIP degradation when we combined both peptide antibodies (Figure 1H, lanes 8-10). In conclusion, Trim-Away is an efficient and specific approach to deplete proteins of interest from various types of *Xenopus* egg extracts.

### Rescue of protein function after cell-free Trim-Away

We next asked whether cell-free Trim-Away can be employed to functionally deplete a protein and whether such depletion can be rescued. To test this, we targeted two genome maintenance mechanisms in egg extract.

Nonhomologous end joining (NHEJ), the major pathway thar repairs DNA double-stranded breaks, directly ligates DNA ends (Stinson and Loparo, 2021). DNA breaks lacking 5’ phosphates require processing by the kinase PNKP prior to repair (Figure 2A). To measure PNKP-dependent NHEJ, we monitored the conversion of a linear DNA molecule lacking 5’ phosphates into open-circular and supercoiled products (Figure 2B, lanes 1-4) (Stinson et al., 2020). Egg extract treatment with only TRIM21 or PNKP antibody did not impair NHEJ, whereas Trim-Away of PNKP in the presence of both components (Figure 2C, lane 11) inhibited end-joining (Figure 2B, lanes 13-16). Importantly, NHEJ was fully restored by the addition of recombinant PNKP to endogenous levels (Figure 2B, lanes 17-20; Figure 2C, lane 12). We ascribe successful rescue to the fact that we used the minimal concentration of TRIM21 needed, and that TRIM21 was partially destroyed during Trim-Away (Figure 2C, compare lanes 9 and 11) so that the added recombinant PNKP was inefficiently targeted (see Discussion).

**Figure 2.**
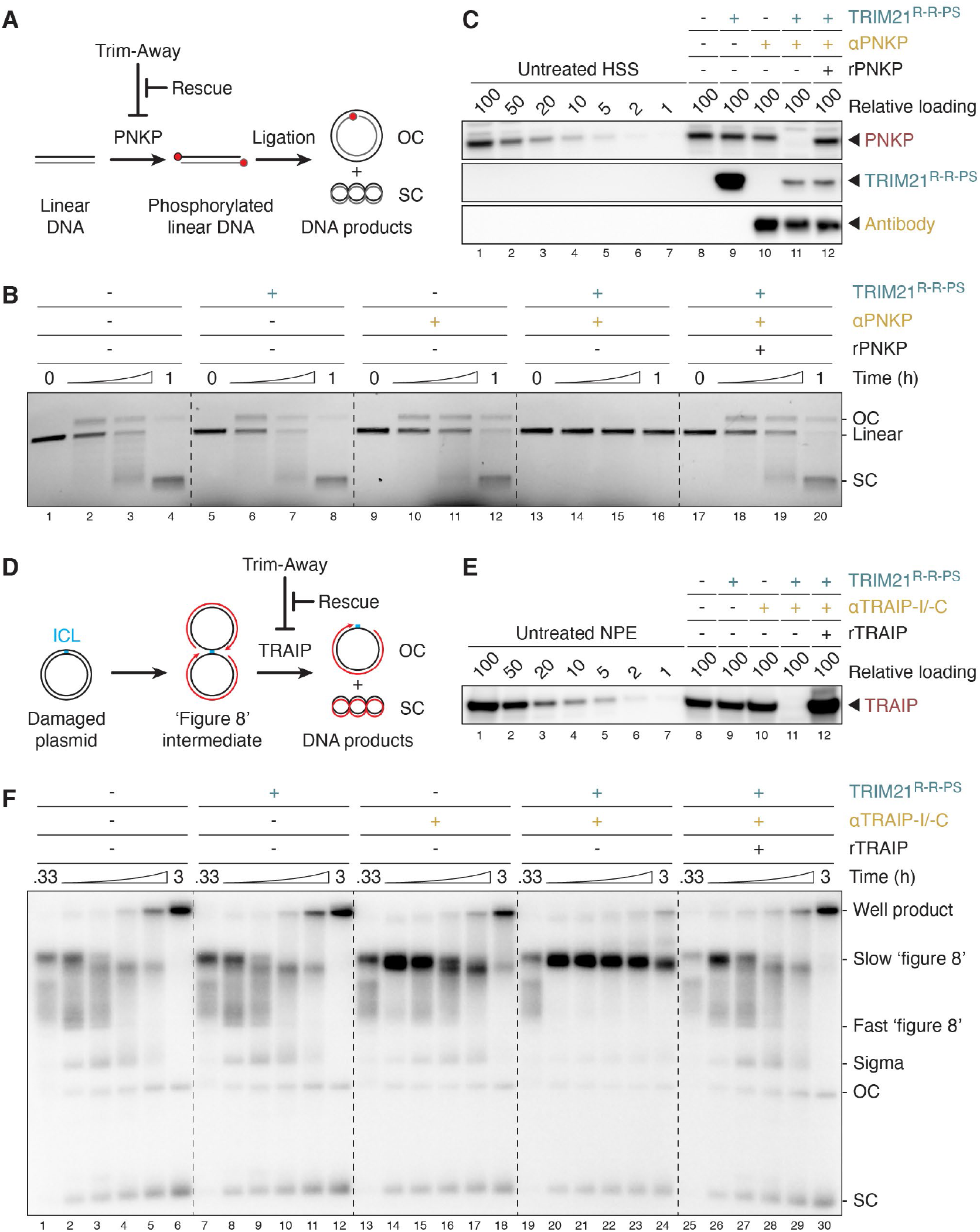
Rescue of protein function after cell-free Trim-Away. **(A)** Schematic of PNKP function during nonhomologous end joining (NHEJ) and experimental setup. Unphosphorylated linear DNA requires PNKP-dependent 5’-hydroxyl phosphorylation (red dot) prior to ligation, forming open-circular (OC) and supercoiled (SC) DNA products. **(B)** NHEJ assay of radiolabeled linear DNA incubated for indicated times with HSS after Trim-Away. Samples were resolved on a native agarose gel and visualized by autoradiography. **(C)** Western blot analysis of HSS treated with TRIM21^R-R-PS^ (final concentration of 375 nM) and PNKP antibody (187.5 nM) for 2 h. In lane 12, recombinant PNKP was supplemented (50 nM) after Trim-Away to rescue the PNKP depletion. **(D)** Schematic of DNA interstrand cross-link (ICL) repair and experimental setup. DNA replication (nascent DNA strands are shown in red) of a plasmid containing a single ICL results in the formation of so-called ‘figure 8’ intermediates. The E3 ubiquitin ligase TRAIP is required for ICL repair, which can be followed by the generation of open-circular (OC) and supercoiled (SC) DNA products. **(E)** Western blot analysis of NPE incubated with TRIM21^R-R-PS^ (500 nM) and two TRAIP peptide antibodies (TRAIP-I and TRAIP-C, 125 nM each) for 2 h. Trim-Away in HSS was performed analogously. After treatment, NPE was supplemented with recombinant TRAIP expressed in wheat germ extract (lane 12) to rescue ICL processing. **(F)** ICL-containing plasmids were replicated in the presence of [*α*-^32^P]dATP, resolved on a native agarose gel and visualized by autoradiography.

Replication-coupled repair of a plasmid containing a DNA interstrand cross-link (ICL) in *Xenopus* egg extracts is initiated when two replisomes converge on the ICL (Figure 2D) (Räschle et al., 2008; Semlow and Walter, 2021). The resulting ‘figure 8’ structure undergoes processing for repair only after replisomes have been unloaded, a reaction that involves CMG helicase ubiquitination by the E3 ubiquitin ligase TRAIP (Wu et al., 2019). When we depleted TRAIP from egg extracts using Trim-Away (Figure 2E, lane 11) and monitored DNA replication by the incorporation of radiolabeled dNTPs, ‘figure 8’ intermediates persisted throughout the entire time course (Figure 2F, lanes 19-24), consistent with functional TRAIP depletion and defective ICL repair. TRIM21^R-R-PS^ alone had no impact on ICL processing, whereas TRAIP antibodies slightly delayed repair, indicating partial TRAIP inhibition by the antibodies themselves (Figure 2F, lanes 1-18). As observed for PNKP, recombinant TRAIP fully restored the functional defect after successful TRAIP destruction (Figure 2E, lane 12; Figure 2F, lanes 25-30). We conclude that cell-free Trim-Away is a powerful approach to interrogate protein function in the *Xenopus* system.

### Target proteins are directly ubiquitinated prior to degradation

According to the current model, the polyubiquitin chain assembled on TRIM21’s N-terminus triggers the proteasomal degradation of TRIM21, antibody, and the target protein, without the latter two components being directly modified (Figure 1B). However, while establishing cellfree Trim-Away, we observed not only modification of TRIM21, but also of antibody and TFIIS (Figure S1B). These modifications were seen in all three egg extracts (Figure S1C), and they were abolished with an E1 inhibitor, demonstrating they reflect ubiquitination (Figure S1D). Since TRIM21 and antibody were added in excess over the endogenous target protein, only a fraction of both were modified. When we performed a time course of Trim-Away instead of the endpoint assay shown in Figure 1G, we detected polyubiquitination of every substrate prior to its degradation (Figures 3 and S2 for darker exposures; Figure S1E). Generally, we observed modification of virtually every protein that was successfully targeted by cell-free Trim-Away, suggesting that direct substrate ubiquitination might be an integral part of the Trim-Away mechanism.

**Figure 3.**
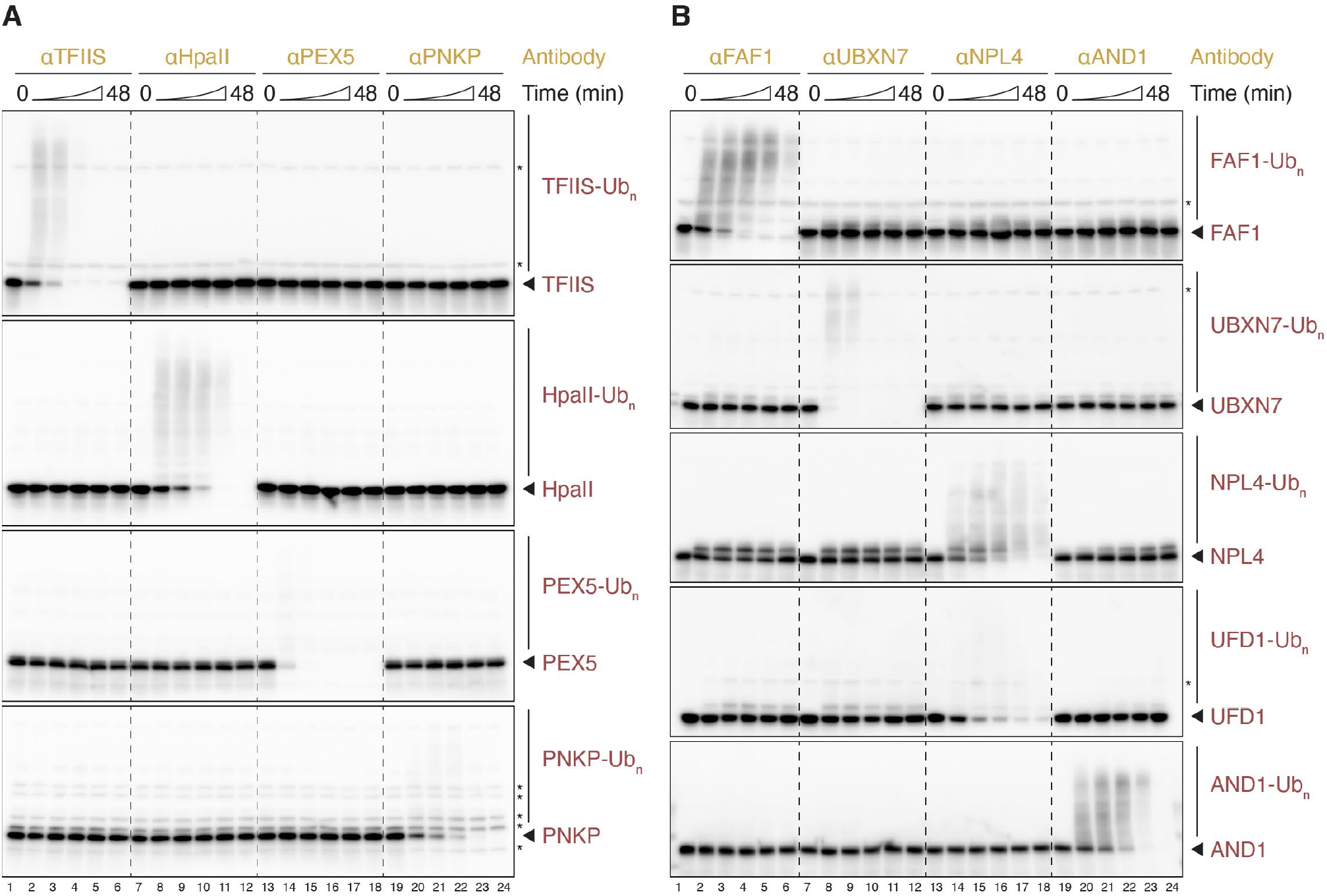
Target proteins are directly ubiquitinated prior to degradation. **(A, B)** The fate of target proteins during cell-free Trim-Away prior to proteasomal degradation was assessed in HSS using TRIM21^R-R-PS^. Comparison of the same endogenous and exogenous substrates targeted in Figure 1G. Samples were taken over a time course (0, 3, 6, 12, 24, and 48 min) prior to SDS-PAGE and western blot analysis. Note that UFD1 is a constitutive binding partner of the targeted protein NPL4. Unmodified proteins of interest are highlighted (arrow). We assume that any bands already present at the start of the reaction are non-specific (asterisk). Longer exposures of the same western blots are shown in Figure S2.

### Direct polyubiquitination of targets is essential for their destruction

To examine whether substrate ubiquitination is required for Trim-Away, we chemically methylated all primary amines in the bacterial DNA binding protein LacI, which blocks its ubiquitination (Figures 4A and S3A). As seen for the proteins targeted in Figure 3, unmethylated LacI was degraded by TRIM21^R-R-PS^ (Figure 4B, lanes 1-6). In contrast, methylation impaired both LacI polyubiquitination and destruction (Figure 4B, lanes 7-12). While this result suggests that LacI ubiquitination is required for Trim-Away, a potential alternative interpretation was that methylation inhibited antibody binding to LacI. Interestingly, unmethylated LacI stimulated TRIM21^R-R-PS^ and antibody ubiquitination (Figure 4B, compare lanes 1-6 and 13-18), consistent with the idea that multiple antibodies assembled on LacI. Importantly, methylated LacI stimulated TRIM21^R-R-PS^ and antibody ubiquitination to the same extent (Figure 4B, lanes 7-12), strongly implying that LacI methylation did not impair antibody recognition, as judged also from comparable immunoblot detection of methylated and unmethylated LacI (Figure 4B, compare lanes 1 and 7). Consistent with antibody ubiquitination, we observed the appearance of an IgG degradation fragment, which was also unaffected by LacI methylation (Figure 4B, “degradation fragment”). Incubation with TRIM21^FL^ for up to 3 h reduced unmethylated LacI levels (Figure 4C), although less efficiently than TRIM21^R-R-PS^. TRIM21^FL^ also failed to destroy methylated LacI, whereas its activation was preserved, as judged from antibody and TRIM21 ubiquitination. Methylation of HpaII had a similar inhibitory effect on Trim-Away by TRIM21^R-R-PS^ and TRIM21^FL^ (Figures S3B and S3C). As an alternative means of inhibiting ubiquitination, we used lysine-less (ΔK) mutants, in which all lysines were replaced by arginine (Figure 4D). Similar to the methylated target proteins, lysine-less PEX5 and TFIIS were neither ubiquitinated nor degraded, whereas TRIM21 and antibody ubiquitination were comparable to wild-type PEX5 and TFIIS, respectively (Figures 4E, 4F, S3D, and S3E). Unlike methylated proteins, lysine-less mutants contain intact N-termini, which implies that cell-free Trim-Away substrates undergo canonical ubiquitination on internal lysines rather than their N-termini. Together, our data reveal that direct target ubiquitination is essential for TRIM21-mediated protein destruction.

**Figure 4.**
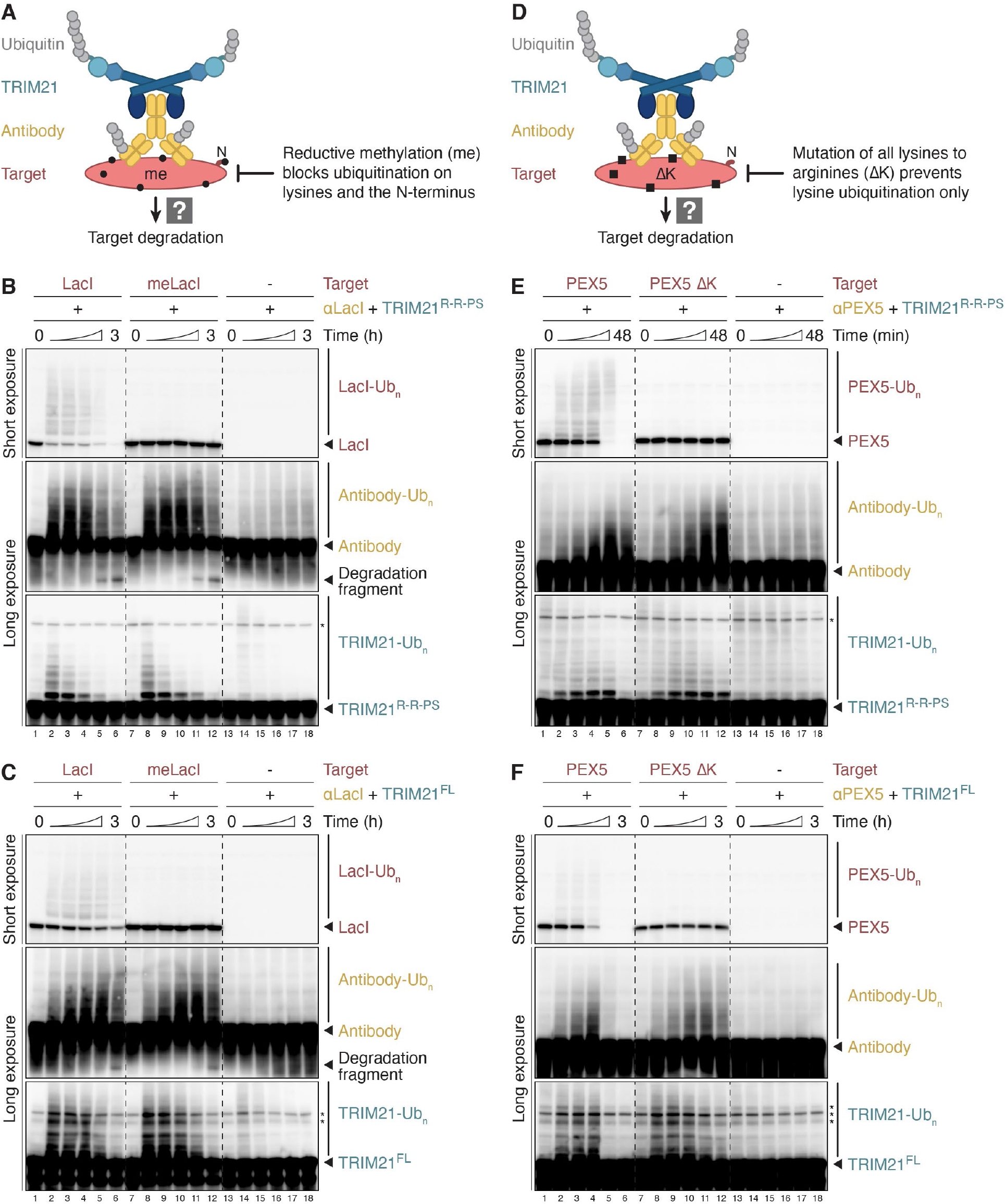
Direct polyubiquitination of targets is essential for their destruction. **(A)** Schematic of experimental setup. To prevent direct target ubiquitination, proteins were chemically methylated in vitro prior to Trim-Away in egg extract. Methylation (black circle) modifies both lysine residues and the N-terminus. **(B, C)** Trim-Away of recombinant LacI variants (final concentration of 250 nM) by TRIM21^R-R-PS^ (B) or TRIM21^FL^ (C) in HSS. IgG degradation fragments are highlighted. **(D)** Schematic of experimental setup. As an alternative means of inhibiting ubiquitination, all lysines (black squares) in target proteins were mutated to arginine (ΔK). Note that lysine-less mutants contain intact N-termini. **(E, F)** Trim-Away of recombinant PEX5 variants (final concentration of 500 nM) by TRIM21^R-R-PS^ (E) or TRIM21^FL^ (F) in HSS. To deplete endogenous PEX5, TRIM21 and antibody were added to egg extract for 15 min (E) or 30 min (F) prior to the addition of recombinant *X. laevis* PEX5 to start cell-free Trim-Away at 0 min. Trim-Away with TRIM21^FL^ required an extended time course (0, 11, 22, 45, 90, and 180 min) for efficient PEX5 degradation.

### Direct target polyubiquitination is sufficient for Trim-Away

To test the current model that TRIM21 ubiquitination is essential for substrate proteolysis, we next methylated TRIM21 as well as the antibody used for Trim-Away (Figures 5A and S4A). As shown in Figure 5B, methylated antibody was just as effective as unmethylated antibody in promoting HpaII degradation (lanes 1-12). Strikingly, methylated TRIM21^R-R-PS^ also supported normal target ubiquitination and destruction, even though TRIM21 itself was no longer ubiquitinated (Figure 5B, lanes 13-18). Similarly, the combination of both methylated antibody and methylated TRIM21^R-R-PS^ did not impair efficient HpaII degradation (Figure 5B, lanes 19-24). We observed similar results with TRIM21^FL^ in an extended time course (Figure 5C). TFIIS was also fully degraded in the presence of methylated antibody and TRIM21, but its Trim-Away kinetics were slightly delayed (Figures S4B and S4C). In this setting, TFIIS polyubiquitination was detectably increased. We speculate that in the absence of antibody and TRIM21 modification, a more complex chain architecture was formed on the substrate, which delayed proteasomal degradation (Figure S4B, lanes 19-24). These data show that among the three key components of Trim-Away, direct ubiquitination of the substrate is necessary and sufficient to support its proteolysis.

**Figure 5.**
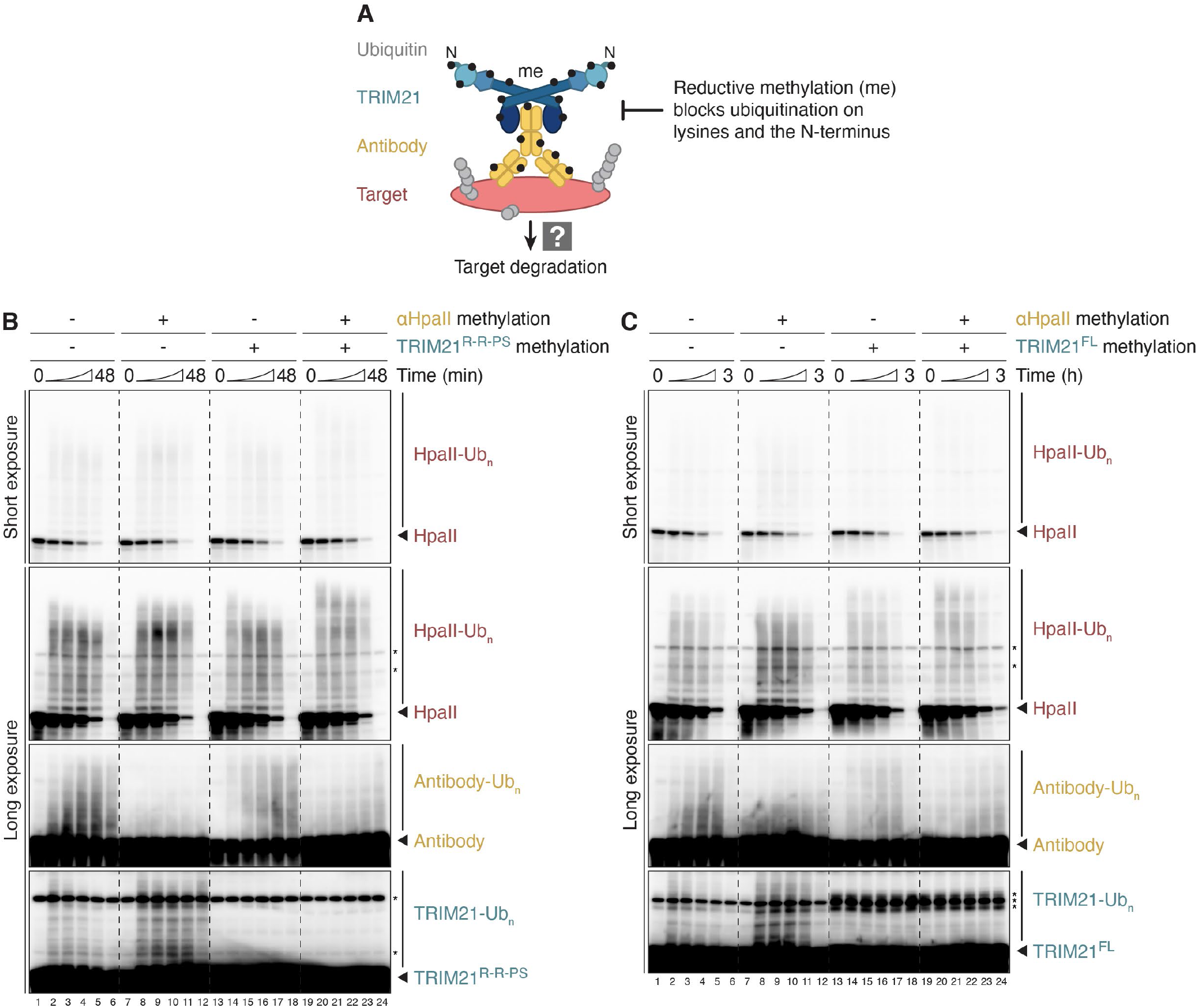
Direct target polyubiquitination is sufficient for Trim-Away. **(A)** Schematic of experimental setup. To prevent direct ubiquitination of TRIM21 or antibody, reductive methylation (black circle) was performed in vitro prior to Trim-Away in egg extract. Note that this procedure inhibits ubiquitination of both the N-terminus and lysine residues. **(B, C)** Cell-free Trim-Away of recombinant unmethylated HpaII (final concentration of 250 nM) by HpaII antibody and TRIM21^R-R-PS^ (B) or TRIM21^FL^ (C). The methylation state of HpaII antibody and TRIM21 variant is indicated. Trim-Away with TRIM21^FL^ required an extended time course (0, 11, 22, 45, 90, and 180 min) for efficient HpaII degradation. Note that methylation of TRIM21^FL^ resulted in an even stronger non-specific band (asterisks), which is already present at the beginning of the Trim-Away reaction.

### Complex heterotypic K48/K63 ubiquitin chains are assembled on Trim-Away targets

Given the importance of direct target polyubiquitination during cell-free Trim-Away, we wanted to characterize the ubiquitin chain architecture involved. To this end, we combined Ubiquitin Chain Restriction (UbiCRest) analysis (Hospenthal et al., 2015; Mevissen et al., 2013) and ubiquitin linkage-specific antibodies. Biotinylated targets were immobilized on streptavidin-coupled magnetic beads and added to egg extract for Trim-Away in the presence of proteasome inhibitor (Figure S5A). After the reaction, the beads were washed and subjected to deubiquitinase (DUB) treatment and western blot analysis. As seen for free HpaII (Figure 3A), immobilized HpaII was efficiently polyubiquitinated only in the presence of TRIM21^R-R-PS^ and HpaII antibody (Figure 6A, lanes 1-4). This high-molecular weight (HMW) signal was not only observed when we blotted with antibodies against HpaII and ubiquitin, but also with antibodies detecting K48- and K63-linked polyubiquitin chains, respectively (Figures 6A-6D, lane 4). The non-specific DUB USP2 converted this HMW signal to monoubiquitin and regenerated unmodified HpaII (Figures 6A and 6B, lane 5). Consistent with the detection of both K48 and K63 chains, the linkage-specific DUBs OTUB1* (cleaves only K48 linkages) and AMSH* (K63 specific) shortened the HMW smear of HpaII (Figure 6A, lanes 6-7). Treatment with both DUBs together resulted in further shortening; since both enzymes are not expected to cleave the proximal ubiquitin that is directly attached to the substrate (Mevissen and Komander, 2017), the remaining modified HpaII species may contain multiple monoubiquitin moieties conjugated to multiple lysine residues on the target, assuming no other linkage types were present (Figure 6A, lane 8).

**Figure 6.**
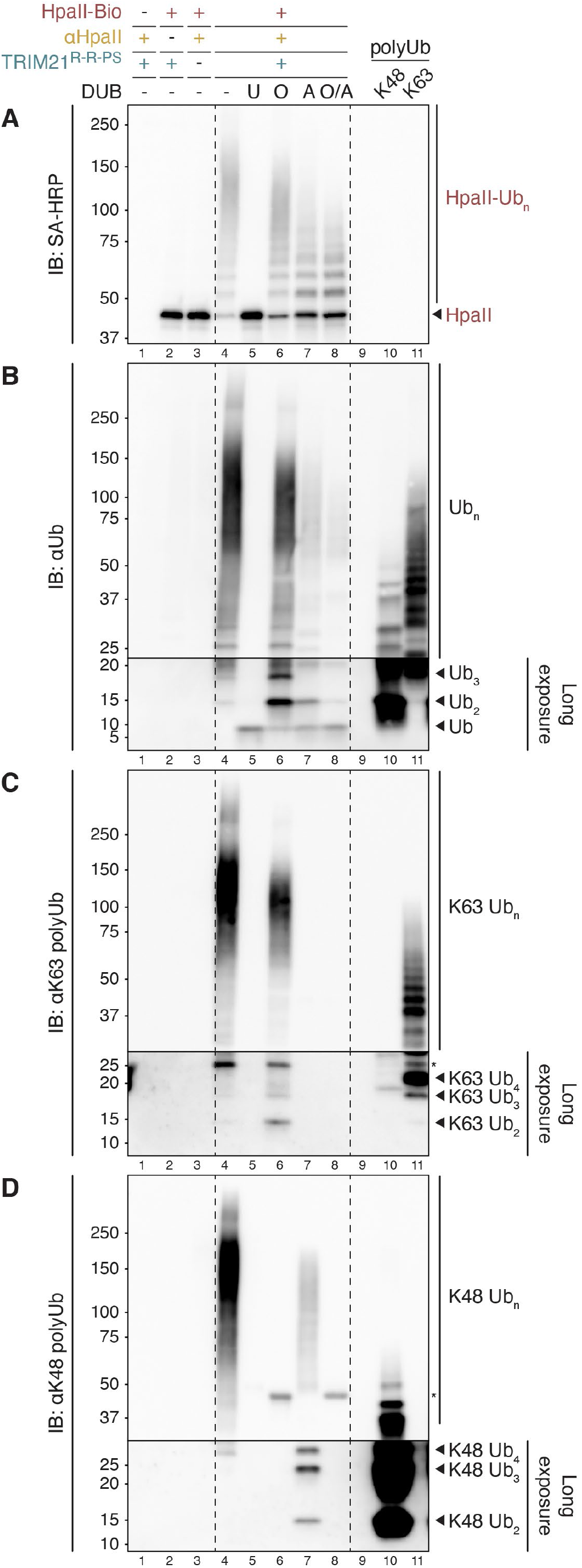
Complex heterotypic K48/K63 ubiquitin chains are assembled on Trim-Away targets. **(A-D)** Ubiquitin Chain Restriction (UbiCRest) analysis of biotinylated HpaII after cell-free Trim-Away (see experimental setup in Figure S5A). The target protein was immobilized on streptavidin-coupled magnetic beads prior to Trim-Away in HSS containing proteasome inhibitor (MG-262). After Trim-Away for 20 min, the beads were washed and subjected to deubiquitinase (DUB) treatment, comparing nonspecific <u>U</u>SP2, K48 linkage-specific <u>O</u>TUB1*, K63specific <u>A</u>MSH*, or both <u>O</u>TUB1* and <u>A</u>MSH*. Digested samples were analyzed by SDS-PAGE and western blotting, visualizing biotinylated HpaII (A), total ubiquitin (B), K63-linked ubiquitin chains (C), or K48-linked ubiquitin chains (D). In lanes 10 and 11, K48- and K63-linked ubiquitin chains assembled in vitro were separated alongside to confirm the specificities of ubiquitin antibodies and for comparing released polyubiquitin species after DUB treatment. The low-molecular weight sections in panels B-D were exposed longer than the high-molecular weight regions to support the identification of unattached ubiquitin species. Asterisks correspond to crossreacting bands.

Treatment with the K63-specific AMSH* removed the majority of total ubiquitin while hydrolyzing all K63 linkages (Figures 6B and 6C, lane 7). Interestingly, AMSH* also released the vast majority of K48 signal from HpaII in the form of unattached K48-linked di-, tri-, and tetraubiquitin, demonstrating that the bulk of K48 linkages was built on top of K63 chains (Figure 6D, lane 7). Despite only slightly reducing the total ubiquitin signal (Figure 6B, lane 6), the K48-specific OTUB1* erased all detectable K48 chains from HpaII, confirming complete treatment (Figure 6D, lane 6). Strikingly, OTUB1* also decreased the K63 signal and released free K63-linked chains, suggesting that a fraction of K63 chains was assembled on K48-linked modifications (Figure 6C, lane 6). In both cases, unattached chains were almost fully hydrolyzed to monoubiquitin when both DUBs were combined, suggesting that only trace amounts of other linkage types were released as well (Figure 6B, lane 8). This analysis was also performed with immobilized LacI, resulting in virtually identical findings (Figures S5B-S5E). Our results reveal a rather complex polyubiquitin chain architecture on Trim-Away substrates. Multiple target lysines appear to be modified, and the predominantly heterotypic K48/K63-mixed or -branched chains lack absolute hierarchy, yet K63 linkages are more favored to be at the base of the chains.

### TRIM21 is ubiquitinated on lysine residues during cell-free Trim-Away

Our data suggest that target proteins are likely polyubiquitinated on multiple lysines prior to their proteasomal degradation (Figures 4 and 6). In contrast, TRIM21 ubiquitination, which is not required for substrate destruction (Figure 5), is proposed to occur on TRIM21’s N-terminus (Fletcher et al., 2015). In addition to triggering protein degradation, N-terminal TRIM21 ubiquitin chains have also been proposed to play a role in activating immune signaling after their release by the proteasome (Fletcher et al., 2015). To better understand potential differences between target and TRIM21 modifications, we examined whether TRIM21 ubiquitination during cell-free Trim-Away occurs on the N-terminus or on lysine side chains (Figure 7A). To this end, we compared the following TRIM21^R-R-PS^ constructs (Figure S6A): i) TRIM21 wildtype (WT), which can be ubiquitinated on both the Nterminus and on lysines, ii) methylated TRIM21 (meWT), which cannot be ubiquitinated on any primary amine, iii) TRIM21 lacking surface-exposed lysines (ΔK), whose N-terminus is available for ubiquitination, and iv) a ubiquitin (K63R)-TRIM21 fusion protein (Ub^K63R^-WT), which can only be ubiquitinated on lysines because UBE2W, which specifically ubiquitinates the N-terminus of proteins, cannot modify the ubiquitin N-terminus (Scaglione et al., 2013; Tatham et al., 2013; Vittal et al., 2015), and because UBE2N/V1-mediated extension of the N-terminal monoubiquitin is prevented by the K63R mutation. In vitro ubiquitination reactions with UBE2W and UBE2N/V1 confirmed that the different TRIM21 constructs contained the expected ubiquitination sites (Figures S7A and S7B).

**Figure 7.**
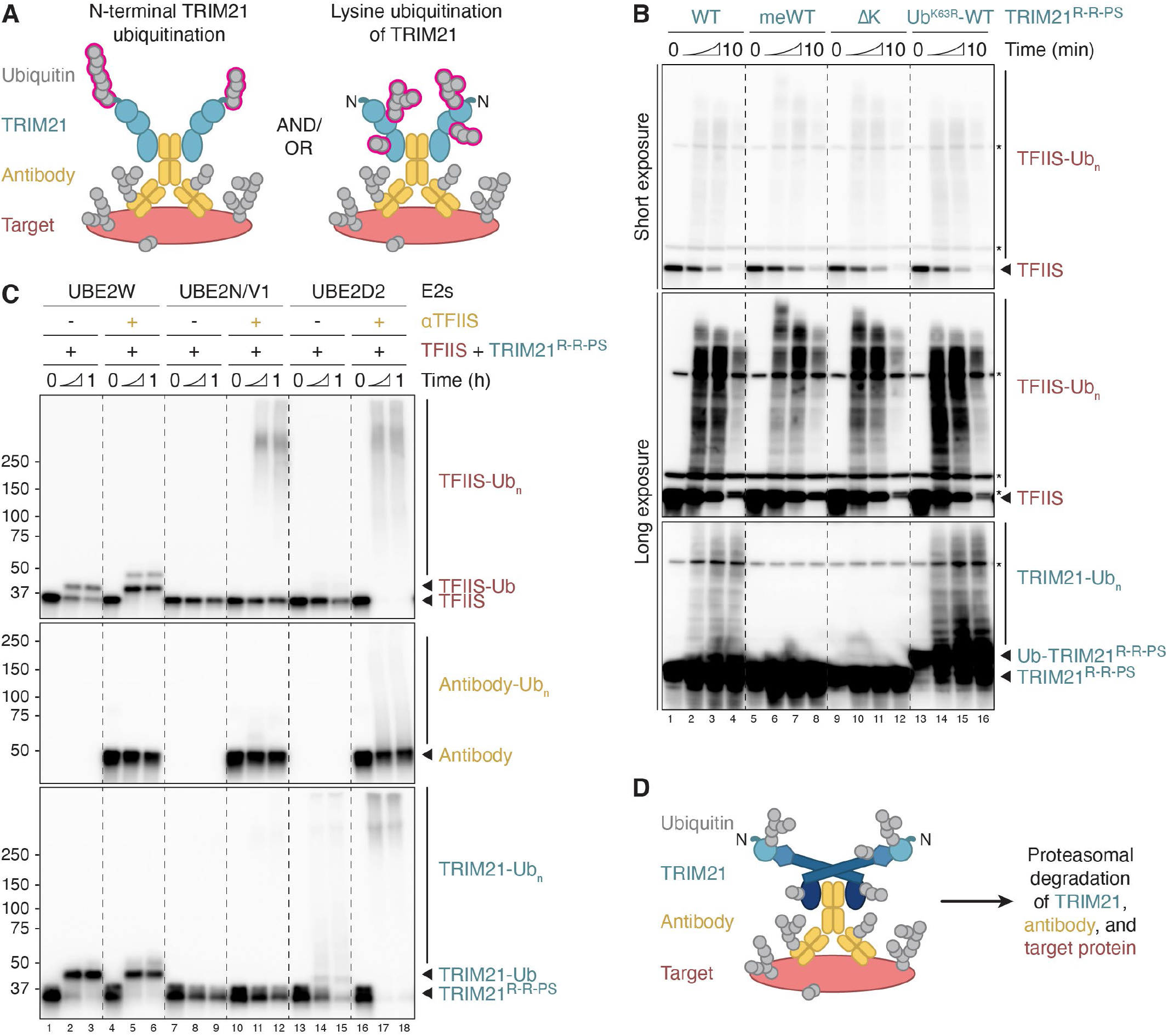
TRIM21 is ubiquitinated on lysine residues during cell-free Trim-Away. **(A)** Schematic comparing potential TRIM21 ubiquitination sites (pink): N-terminus (left panel) and/or lysine residues (right panel). **(B)** Cell-free Trim-Away assay targeting endogenous TFIIS in HSS. Comparison of the following TRIM21^R-R-PS^ constructs: i) Wild-type (WT), which can be modified on both the N-terminus and on lysines, ii) methylated TRIM21 (meWT), which cannot be ubiquitinated on any primary amine, iii) lysine-less TRIM21 (ΔK) containing a ubiquitinatable N-terminus, and iv) a ubiquitin (K63R)-TRIM21 fusion protein (Ub^K63R^-WT), which can be ubiquitinated on lysines only. See Figure S6 for additional schematics. **(C)** Reconstitution of antibody-dependent target ubiquitination by TRIM21. Recombinant TFIIS was incubated with ATP, ubiquitin, E1, indicated E2 enzymes, and TRIM21^R-R-PS^ in the absence or presence of TFIIS antibody. **(D)** Proposed Trim-Away model. Upon TRIM21 recruitment to antibody-bound targets, E3 ligase activation leads to direct polyubiquitination of all components on lysines. Substrate modification with heterotypic K48/K63-mixed or branched polyubiquitin chains mediates its proteasomal degradation.

Trim-Away in egg extract revealed that all four TRIM21 constructs efficiently mediated the degradation of endogenous TFIIS (Figure 7B). As expected and observed above (Figures 5B and S4B), methylated TRIM21^R-R-PS^ was not ubiquitinated during cell-free Trim-Away (Figure 7B, lanes 5-8). Importantly, TRIM21^R-R-PS^ ΔK was also not modified, whereas Ub^K63R^-TRIM21^R-R-PS^ featuring a blocked N-terminus was readily polyubiquitinated (Figure 7B, lanes 9-16). Target ubiquitination was again enhanced when TRIM21 ubiquitination was absent (Figure 7B, lanes 5-12). In summary, these data clearly demonstrate that all of the observed TRIM21 polyubiquitination during cell-free Trim-Away occurs on lysine side chains and not on its N-terminus (Figure S6B), as seen also for substrate ubiquitination (Figure 4).

### Reconstitution of antibody-dependent target ubiquitination by TRIM21

The model that UBE2W first monoubiquitinates the Nterminus of TRIM21, followed by UBE2N/V2-mediated extension with K63-linked chains, is largely based on reconstituted biochemical reactions (Fletcher et al., 2015; Kiss et al., 2019, 2021). We confirmed that this sequence of events can occur using TRIM21^R-R-PS^ constructs (Figure S7A and S7B). However, previous studies focused mainly on these E2s and monitored TRIM21 autoubiquitination in the absence of antibody and substrate. In addition to UBE2W, UBE2N/V1 and UBE2N/V2, TRIM21 also operates in conjunction with UBE2D family members (Kiss et al., 2019). Accordingly, TRIM21^R-R-PS^ assembled free ubiquitin chains in the presence of UBE2D2, as seen for UBE2N/V1 (Figure S7C). In contrast to UBE2N/V1, however, UBE2D2mediated TRIM21 autoubiquitination occurred even without UBE2W, indicating lysine modification (Figure S7C, lanes 13-18). We next wanted to reconstitute antibody-dependent TRIM21 auto- and substrate polyubiquitination with purified proteins, which could shed light on which E2s participate in cell-free Trim-Away. In the presence of UBE2W, both TRIM21^R-R-PS^ and the target TFIIS were readily monoubiquitinated, even without antibody (Figure 7C, lanes 1-6). In contrast, UBE2N/V1 promoted antibody-dependent TFIIS polyubiquitination, but unlike cell-free Trim-Away, antibody and TRIM21^R-R-PS^ were not modified (Figure 7C, lanes 7-12). Strikingly, in the presence of UBE2D2, the addition of antibody stimulated not only efficient ubiquitination of TFIIS, but also of TRIM21^R-R-PS^ and antibody (Figure 7C, lanes 13-18). Comparable results were obtained with TRIM21^FL^ (Figure S7D). UbiCRest analysis of polyubiquitin chains assembled with TRIM21^FL^ and UBE2D2 in vitro identified heterotypic K48/K63 linkages (Figure S7E). Finally, reconstituted antibody-dependent LacI and TFIIS modification by TRIM21^FL^ and UBE2D2 revealed polyubiquitination patterns closely resembling those observed in egg extract (Figures S7F and S7G, compare to Figures 6A and S5B). In summary, we have reconstituted the ubiquitination events underlying cell-free Trim-Away. Consistent with our ubiquitin linkage type characterization in egg extract, this analysis suggests that UBE2D family members and possibly UBE2N/V1, but unlikely UBE2W, underlie this targeted protein degradation pathway.

## DISCUSSION

The F_c_ receptor TRIM21 is an E3 ubiquitin ligase with two crucial roles after antibody-decorated pathogens enter the cytoplasm: i) activating immune signaling, and ii) triggering proteasome-mediated pathogen destruction (Mallery et al., 2010; McEwan et al., 2013). The latter activity has been exploited for targeted protein degradation in Trim-Away (Clift et al., 2017). Here, we have established cell-free Trim-Away, which we show operates by a mechanism that is fundamentally different from the one previously proposed.

### Cell-free Trim-Away – a novel tool to interrogate protein function

Trim-Away was initially developed as a powerful approach for the acute degradation of endogenous proteins in a wide range of cell types and embryos, including *Xenopus laevis* (Chen et al., 2019; Clift et al., 2017; Israel et al., 2019; Weir et al., 2021). Here, we demonstrate rapid and efficient Trim-Away in cell-free extracts derived from *Xenopus* eggs (Figure 1). In contrast to cell-based systems, where the delivery of Trim-Away components can be technically challenging, extract systems are highly tractable. Upon addition of recombinant human TRIM21 and antibody to egg extract, target degradation kinetics depend largely on substrate abundance, and we observed complete degradation of multiple proteins in less than 12 minutes (Figures 3 and S2; TFIIS, PEX5, and UBXN7). In contrast, immunodepletions generally take hours and require at least 50-fold more antibody than cell-free Trim-Away. In principle, extracts can be supplemented with TRIM21 and antibodies at any time, even mid-way through a reaction to test whether a protein functions at multiple steps of a complex biological process. Another potential application for cell-free Trim-Away is the targeting of transmembrane proteins, whose immunodepletion would inadvertently also remove the surrounding structure, such as an entire organelle.

Finally, the fate of target proteins can be monitored with unprecedented temporal resolution, which helped us uncover novel features of antibody-mediated target degradation by TRIM21 (see below). Although cell-free Trim-Away is highly specific (Figure 1G), we detected UFD1 ubiquitination and degradation upon NPL4 antibody treatment (Figure 3B, lanes 13-18), indicating that stable binding partners of targeted proteins may be codepleted.

To achieve efficient cell-free Trim-Away, extracts need to be supplemented with antibody and TRIM21 in excess over the target protein (Figure 1C). We show that the minimal TRIM21^R-R-PS^ construct featuring the PRYSPRY domain fused to two RING domains (Kiss et al., 2021) promotes more efficient cell-free Trim-Away than full-length TRIM21^FL^, which required longer incubation times. Targeting monomeric proteins with monomeric antibodies is generally not successful due to the lack of target-induced TRIM21 clustering (Zeng et al., 2021). While testing a large panel of antibodies (Figures 1G and 1H), we noticed that polyclonal antibodies raised against entire proteins or domains were much more likely to promote successful Trim-Away compared to peptide antibodies. We speculate that the small epitopes of peptide antibodies generally allow only a very limited number of antibodies to bind to the substrate, which may not recruit sufficient TRIM21 molecules for target ubiquitination, even when TRIM21^R-R-PS^ is used. To circumvent this problem, we combined two peptide antibodies raised against an internal and a C-terminal region of TRAIP to promote efficient depletion, which was not observed with either peptide antibody alone (Figure 1H).

Using two well-established genome maintenance pathways, we demonstrate full functional rescues after cell-free Trim-Away (Figure 2), which is crucial to rule out off target effects and to allow structure-function analyses. To achieve rescue, it was important to determine the lowest amount of TRIM21 and antibody that yields target depletion. This is because a fraction of TRIM21 and antibody gets ubiquitinated and degraded during Trim-Away (Figure 2C, compare lanes 9-11), such that the remaining TRIM21 and antibody are unable to trigger effective destruction of recombinant proteins added for rescue. Our observations are consistent with Trim-Away phenotypes being rescuable after target overexpression in human cells (Clift et al., 2017).

### The mechanism of target protein degradation by TRIM21

TRIM21 function has been suggested to rely solely on K63-linked polyubiquitination on TRIM21’s N-terminus (Figure 1B) (Fletcher et al., 2015; Kiss et al., 2019; Zeng et al., 2021), while antibody and pathogen (i.e., viral capsid) modifications were not detected (Mallery et al., 2010). Surprisingly, we observed direct polyubiquitination of virtually all antibodies and target proteins in addition to TRIM21 during cell-free Trim-Away (Figures 3 and S2; proposed model in Figure 7D). Antibody and substrate modification have never been comprehensively assessed in cells and may have escaped detection in previous Trim-Away experiments because western blots were performed at single end points (Clift et al., 2017; Kiss et al., 2019, 2021; Zeng et al., 2021). Our cell-free system allowed us to monitor the fate of target proteins prior to degradation. Substrate destruction crucially depended on its direct polyubiquitination on lysine residues (Figure 4), while antibody and TRIM21 modification were dispensable (Figure 5). Our data predict that viruses lacking surface exposed lysines would escape destruction, but still support immune signaling. Direct target ubiquitination in the context of virus neutralization is consistent with the essential role of the p97 ATPase (Hauler et al., 2012), and the recent finding that p97 initiates substrate processing by acting on and unfolding substrate-bound ubiquitin chains (Twomey et al., 2019). In our study, p97 inhibition delayed target proteolysis of even a monomeric substrate (Figure 1D), suggesting that p97-assisted unfolding enhances proteasomal degradation and may play a general role in Trim-Away.

A key question concerns which E2 enzymes collaborate with TRIM21 (Kiss et al., 2019). The sequential interaction with UBE2W and UBE2N/V2 promotes TRIM21’s N-terminal mono- and subsequent K63-linked polyubiquitination in vitro in the absence of antibody or target protein (Fletcher et al., 2015), which we confirmed using TRIM21^R-R-PS^ (Figure S7A and S7B). However, by comparing various TRIM21 constructs in cell-free Trim-Away, we found that TRIM21 was exclusively modified on lysines and not on its N-terminus (Figures 7B and S6). We further tested different E2 enzymes in reconstituted reactions of antibody-mediated substrate ubiquitination by TRIM21 (Figures 7C and S7D). UBE2W efficiently monoubiquitinated both TRIM21 and the substrate TFIIS even in the absence of antibody. It should be noted that in vitro assays in this and previous studies were performed with unphysiologically high UBE2W concentrations (0.5-1 µM). While UBE2N and UBE2Ds are highly abundant in *Xenopus* eggs (UBE2N, 880 nM; UBE2D2, 260 nM; UBE2D4, 710 nM), the UBE2W concentration is more than 100-fold lower (5.8 nM) (Wühr et al., 2014). Overall, there is no evidence to support a role for UBE2W or N-terminal TRIM21 ubiquitination in cell-free Trim-Away, raising the question of whether this concept is relevant in cells. In this regard, it is important to note that N-terminal TRIM21 ubiquitination has never been directly shown in cells, and that UBE2W protein was not detected in U2OS cells (Beck et al., 2011), a cell line that works well for Trim-Away (Clift et al., 2017). Although UBE2W knockdown was shown to impair virus neutralization and immune signaling (Fletcher et al., 2015), this could have been an indirect effect. Thus, the role of N-terminal ubiquitination for cellular TRIM21 function should be revisited.

In addition to UBE2W, members of the UBE2D family as well as UBE2N/V1 and UBE2N/V2 also work in conjunction with TRIM21 (Kiss et al., 2019). Importantly, we detected a robust antibody dependence for UBE2N/V1- and UBE2D2-mediated ubiquitination events by TRIM21 in reconstituted reactions (Figures 7C and S7D). UBE2N is highly K63 ubiquitin linkage specific, whereas the more promiscuous UBE2D family predominantly assembles K48- and K63-linked polyubiquitin chains with different RING E3 ligases in vitro (Wauer et al., 2015), including branched structures (Swatek et al., 2019). Consistent with potential roles for these E2 enzymes in antibody-mediated TRIM21 activity, we show that heterotypic K48- and K63-linked polyubiquitin chains were synthesized on substrates during cell-free Trim-Away (Figures 6 and S5). K63 linkages were located preferentially at the base with K48-linked chains built on top, suggesting an architecture similar to the polyubiquitination of overexpressed TRIM21 (Fletcher et al., 2015). Finally, we also detected heterotypic K48/K63 ubiquitin chains in reconstituted reactions with TRIM21^FL^ and UBE2D (Figure S7E), and experiments further including antibody and target proteins closely resembled substrate polyubiquitination patterns observed during cell-free Trim-Away (Figures S7F and S7G). Although knockdown of three of the four UBE2D family members does not impact virus neutralization (Kiss et al., 2019), incomplete protein depletion might explain the lack of a phenotype. In contrast, UBE2N knockdown reduces NF-*κ*B induction and virus neutralization (Fletcher et al., 2015; McEwan et al., 2013), yet these effects were attributed to K63-linked ubiquitin chains that were unattached or assembled on N-terminally monoubiquitinated TRIM21, respectively. We propose that UBE2N-dependent K63 ubiquitination contributes to immune system activation (McEwan et al., 2013), while UBE2Ds may also trigger the destruction of pathogens and Trim-Away targets.

## Conclusion

In summary, our results expand the use of Trim-Away to interrogate protein function in cell-free systems. Moreover, by showing that substrate but not TRIM21 ubiquitination is critical for Trim-Away, they suggest that current models of how TRIM21 promotes antibody-mediated target destruction should be revisited.

## ACKNOWLEDGEMENTS

We thank Tom Rapoport, Sichen Shao, Eric Fischer, Sun Hur, Jonathan Pruneda, Michael Skowyra, Olga Kochenova, and Maksym Shyian for critical feedback on the manuscript. We are also grateful for advice and reagents from Tom Rapoport and Michael Skowyra (recombinant PEX5 variants and PEX5 antibody), Leo Kiss and Leo James (previously unpublished TRIM21^R-R-PS^ sequence), Julien Duxin (purified LacI antibody), Ben Stinson (NHEJ substrate and recombinant PNKP), Alex Wu (pICL and TRAIP antibodies), Gheorghe Chistol and Emily Low (purified AND1 antibody), Justin Sparks (recombinant M.HpaII-Bio variants), Geylani Can (methylated LacI), and Yang Lim (LSS). T.E.T.M. was supported by an HHMI fellowship of The Jane Coffin Childs Memorial Fund for Medical Research. A.V.P. was supported by the BCMP Summer Scholars Program. This work was supported by NIH grant HL098316. J.C.W. is a Howard Hughes Medical Institute Investigator and an American Cancer Society Research Professor.

## AUTHOR CONTRIBUTIONS

T.E.T.M. and A.V.P. performed all experiments. T.E.T.M. and J.C.W. conceived experiments, and analyzed and wrote the manuscript with input from A.V.P.

## METHODS

### *Xenopus laevis* and egg extracts

Experiments involving adult female (Nasco #LM0053MX) and male (Nasco #LM00715MX) *Xenopus laevis* were approved by the Harvard Medical Area Institutional Animal Care and Use Committee and conform to all relevant regulatory standards. Egg extracts (LSS, low-speed supernatant; HSS, high-speed supernatant; and NPE, nucleoplasmic extract) were prepared as described previously (Gillespie et al., 2012; Sparks and Walter, 2019).

### Protein expression and purification

The expression plasmid for full-length human TRIM21 (HLTV-hTRIM21, referred to as TRIM21^FL^) was ordered from Addgene (#104973). Recombinant TRIM21^FL^ was expressed and purified as described previously (Clift et al., 2018). AMSH* (Addgene #66712) and OTUB1* (Addgene #65441) were produced according to (Michel et al., 2015). Biotinylated LacI and M.HpaII (referred to as HpaII) were expressed and purified as described previously (Duxin et al., 2014).

The sequence encoding the minimal TRIM21 construct containing the PRYSPRY domain fused to two RING domains (TRIM21^R-R-PS^) was received from Leo Kiss and ordered as a codon-optimized gene block from Integrated DNA Technologies (IDT). Using Gibson Assembly (NEBuilder® HiFi DNA Assembly Master Mix, New England Biolabs), the gene block was directly cloned into the pOPINK vector (Addgene #41143), featuring a PreScission protease cleavable N-terminal His6-GST tag. Expression plasmids for TRIM21^R-R-PS^ ubiquitin fusion constructs (Ub^WT^-TRIM21^R-R-PS^ and Ub^K63R^-TRIM21^R-R-PS^; both constructs contain additional Ub G75/76A mutations to prevent DUB cleavage), TRIM21^R-R-PS^ lacking surface-exposed lysines (ΔK; this construct contains a single lysine that is buried within the PRYSPRY domain), as well as *X. laevis* TFIIS wild-type and ΔK were assembled analogously. All plasmids were verified by Sanger sequencing.

All TRIM21^R-R-PS^ variants were expressed in *E. coli* OverExpress™ C41(DE3) Chemically Competent Cells (Sigma #CMC0017) grown at 37°C in LB media supplemented with the appropriate antibiotics. After reaching an OD_600_ of 0.5-0.7, cells were induced with 0.5 mM IPTG, supplemented with 20 µM ZnSO_4_, and grown for 16-20 h at 18°C. Protein purifications were performed at 4°C. Bacterial cells from 2 l were lysed by sonication in 40 ml TRIM21 lysis buffer (25 mM HEPES [pH 8.0], 300 mM NaCl, 20 µM ZnSO_4_, 10% glycerol, and 2 mM DTT) supplemented with one tablet cOmplete™ EDTA-free Protease Inhibitor Cocktail (Roche) and 1 mg/ml lysozyme, and cleared by centrifugation at 35,000 rpm in a Beckman Ti45 rotor for 1 h. The clarified lysate was incubated with 2 ml pre-equilibrated Glutathione Sepharose 4B resin (Cytiva) for 1 h at 4°C on a rotating wheel. The resin was washed extensively with TRIM21 wash buffer (25 mM HEPES [pH 8.0], 500 mM NaCl, 10 µM ZnSO_4_, 5% glycerol, and 2 mM DTT) and equilibrated with TRIM21 gel filtration buffer (25 mM HEPES [pH 8.0], 200 mM NaCl, 10 µM ZnSO_4_, and 2 mM DTT). The Nterminal His6-GST tag was cleaved on the resin with GST-tagged PreScission protease overnight at 4°C on a rotating wheel. The released protein was collected in 5-10 ml gel filtration buffer, concentrated to 2 ml using an Amicon 10 MWCO spin concentrator (Millipore), and subjected to size-exclusion chromatography on a HiLoad® 16/600 Superdex® 75 pg column (Cytiva) equilibrated in gel filtration buffer. Pooled peak fractions were concentrated, snap-frozen in liquid nitrogen, and stored at −80°C. Purified TRIM21^R-R-PS^ variants are shown Figure S1A.

*X. laevis* TFIIS variants were expressed and purified as described above, except for using 0.2 mM IPTG for induction and the following purification buffers: TFIIS lysis buffer (50 mM HEPES [pH 7.5], 500 mM NaCl, 20 µM ZnSO_4_, 10% glycerol, and 2 mM DTT), TFIIS wash buffer (25 mM HEPES [pH 7.5], 500 mM NaCl, 10 µM ZnSO_4_, 5% glycerol, and 2 mM DTT), and TFIIS gel filtration buffer (25 mM HEPES [pH 7.5], 200 mM NaCl, 10 µM ZnSO_4_, 5% glycerol, and 2 mM DTT). Size-exclusion chromatography was performed for *xl*TFIIS, whereas *xl*TFIIS ΔK was directly concentrated, snapfrozen, and stored at −80°C after overnight cleavage with PreScission protease.

### Reductive methylation

HpaII was methylated under denaturing conditions as described previously (Larsen et al., 2019) and kindly provided by Justin Sparks. LacI, antibodies against HpaII and LacI, as well as TRIM21^FL^ and TRIM21^R-R-PS^ were methylated under native conditions using the Reductive Alkylation Kit (Hampton Research, HR2-434). Prior to the procedure, proteins stored in Tris buffer were dialyzed four times against methylation buffer (25 mM HEPES [pH 7.5], 200 mM NaCl, and 10% glycerol) using Slide-ALyzer™ Dialysis Cassettes (Thermo Scientific) with an appropriate cut-off. Reductive methylation reactions were initiated by the addition of the dimethylamine borane complex and formaldehyde according to manufacturer’s instructions. After overnight incubation at 4°C, reactions were stopped with 100 mM Tris [pH 7.5] and 5 mM DTT (final concentrations), except DTT was not added to antibody reactions. Methylated TRIM21 was dialyzed twice against TRIM21 gel filtration buffer (25 mM HEPES [pH 8.0], 200 mM NaCl, 10 µM ZnSO_4_, and 2 mM DTT). Methylated antibodies were dialyzed twice against antibody storage buffer (50 mM Tris [pH 7.5], 150 mM NaCl, and 10% sucrose). Methylated LacI was dialyzed twice against LacI dialysis buffer (50 mM Tris [pH 7.5], 1 mM EDTA, 100 mM NaCl, 1 mM DTT, and 30% glycerol). All proteins were subsequently concentrated using Amicon spin concentrators (Millipore), snap-frozen in liquid nitrogen, and stored at −80°C. Direct comparesons of unmethylated and methylated proteins are shown in Figures S3A and S4A.

### Antibodies and immunodepletion

Rabbit polyclonal antibodies raised against the following proteins were used for both western blotting and Trim-Away unless otherwise indicated: M.HpaII (Larsen et al., 2019); PEX5 (Romano et al., 2019); PNKP (Stinson et al., 2020); FAF1 (Trim-Away only), UBXN7, NPL4 (Trim-Away only) (Kochenova et al., submitted); TRAIP-C (Dewar et al., 2017). The following antibodies used for western blotting are commercially available or were described previously: TRIM21 (Cell Signaling Technology, 92043); FAF1 (Abcam, ab202298); NPL4, UFD1, p97 (Heubes and Stemmann, 2007); Histone H3 (Cell Signaling Technology, 9715); ubiquitin (Santa Cruz Biotechnology, sc-8017); K48-linkage specific polyubiquitin (Cell Signaling Technology, 8081); K63-linkage specific polyubiquitin (Abcam, ab179434).

Rabbit polyclonal TRAIP antibodies (referred to as TRAIP-I; kindly provided by Alex Wu) used for Trim-Away were raised and affinity-purified by Vivitide against a synthetic internal peptide of *X. laevis* TRAIP (amino acids 336-351; Ac-APFKKMKFDNKEHPLSC-amide). Rabbit polyclonal antibodies targeting full-length *X. laevis* TFIIS or an internal fragment of *X. laevis* AND1 (amino acids 846-1127; kindly provided by Gheorghe Chistol and Emily Low) were prepared by Pocono Rabbit Farm and Laboratory and used for western blotting, Trim-Away, and immunodepletion (TFIIS). Rabbit polyclonal LacI antibodies (a generous gift of Julien Duxin) were raised by BioGenes GmbH (Germany) against LacI-Bio and used for both western blotting and Trim-Away. TFIIS, AND1, and LacI antibodies were affinity-purified from serum using the appropriate antigen coupled to AminoLink Coupling Resin (Thermo Scientific) according to manufacturer’s protocol.

For TFIIS immunodepletions (Figures S3D and S3E), one volume of affinity-purified TFIIS antibody (1 mg/ml) was incubated with two volumes of Dynabeads Protein A slurry (Invitrogen) equilibrated with 1x PBS supplemented with 0.25 mg/ml BSA. After overnight incubation on a rotating wheel at 4°C, the resin was washed three times each with 1x PBS supplemented with 0.1 mg/ml BSA and egg lysis buffer (ELB; 10 mM HEPES [pH 7.7], 50 mM KCl, 2.5 mM MgCl_2_, and 250 mM sucrose) supplemented with 0.1 mg/ml BSA. Three volumes of precleared HSS were then depleted with two volumes of antibody-bound beads for three rounds on a rotating wheel for 1 h each at 4°C. The depleted egg extract was collected and immediately used for cell-free Trim-Away as described below.

### Cell-free Trim-Away

After thawing appropriate amounts of egg extract, low-speed supernatant (LSS) and high-speed supernatant (HSS) were supplemented with 3 µg/ml nocodazole (final concentration). ATP regenerating system (2 mM ATP, 20 mM phosphocreatine, and 5 μg/ml phosphokinase; final concentrations) and, where indicated, inhibitors (200 µM final concentration) or exogenous proteins were added to LSS, HSS, and nucleoplasmic extract (NPE) at room temperature for 15 min before starting Trim-Away reactions. HSS and antibodies were centrifuged at 14,000 x *g* for 5 min at room temperature to remove any precipitation. 10x concentrated TRIM21/antibody stock solutions containing TRIM21 (10 µM TRIM21^R-R-PS^ or 25 µM TRIM21^FL^, unless otherwise indicated) and the indicated antibody (5 µM, unless otherwise indicated) were pre-incubated for 10 min at room temperature. Cellfree Trim-Away reactions were started by the addition of nine volumes of egg extract mixture to one volume of 10x TRIM21/antibody stock solution and incubated at room temperature. For SDS-PAGE and western blot analysis, samples were stopped at indicated times as described below. For experiments shown in Figures 1C-1F, samples were withdrawn at indicated times and treated with USP2 (Bio-Techne; 2.5 µM final concentration), E1 inhibitor MLN7243 (Selleck Chemicals; 200 µM), and proteasome inhibitor MG-262 (Bio-Techne; 200 µM) for 30 min at 37°C prior to SDS-PAGE and western blot analysis. For experiments shown in Figures 4B and 4D, cell-free Trim-Away of endogenous PEX5 was started at −15 min as described above. At 0 min, recombinant PEX5 variants were added (500 nM final concentration) to initiate the reactions. For experiments with immobilized target proteins shown in Figures 6 and S5, biotinylated HpaII or LacI were bound to streptavidin-coated magnetic beads (Dynabeads® M-280 Streptavidin) for 30 min at room temperature. Beads were extensively washed with washing buffer (25 mM HEPES [pH 8.0], 200 mM NaCl, 2 mM DTT, 0.25 mg/ml BSA, and 0.02% Tween-20) before Trim-Away components were added to start the reactions. After 20 min, beads were immediately washed with high-salt buffer (50 mM Tris [pH 7.5], 1 M NaCl, 10 mM DTT, and 0.02% Tween-20) prior to Ubiquitin Chain Restriction (UbiCRest) analysis (see below).

### SDS-PAGE and western blotting

At indicated time points, samples were stopped with 2x Laemmli sample buffer (120 mM Tris [pH 6.8], 4% SDS, 20% glycerol, 0.02% bromophenol blue, and 10% βmercaptoethanol), boiled for 2 min at 95°C, and resolved on 4-15% Mini-PROTEAN® TGX™ Precast Protein Gels (Bio-Rad) using Tris-Glycine-SDS Running Buffer (Boston BioProducts). Coomassie staining was performed with InstantBlue® Protein Stain (Expedeon). For western blotting, SDS-PAGE gels were transferred to PVDF membranes (Perkin Elmer). Membranes were blocked in 1x PBST containing 5% (w/v) non-fat milk for 30-60 min at room temperature. Primary antibodies were diluted to 1:300 – 1:10,000 in 1x PBST containing 1% BSA and 0.02% sodium azide and incubated with membranes overnight at 4°C. Membranes were extensively washed with 1x PBST containing 5% (w/v) non-fat milk at room temperature. The following horseradish peroxidase (HRP)-conjugated secondary antibodies were diluted to 1:20,000 in 1x PBST containing 5% (w/v) non-fat milk: Goat Anti-Rabbit IgG (H+L), Mouse Anti-Rabbit IgG (L), and Rabbit Anti-Mouse IgG (H+L) (all Jackson ImmunoResearch). Detection of antibodies after Trim-Away was either performed with the HRP-conjugated Goat Anti-Rabbit IgG (H+L) secondary antibody directly, or by using an unconjugated Mouse Anti-Rabbit IgG (H+L) (Jackson ImmunoResearch) as a primary antibody prior to incubation with the HRP-conjugated Rabbit AntiMouse IgG (H+L) secondary antibody. To detect biotinylated target proteins, blocked membranes were directly incubated with HRP-conjugated streptavidin (Abcam) diluted to 1:2,500 in 1x PBST containing 1.25% (w/v) non-fat milk. All secondary antibody incubations were performed for 45-60 min at room temperature. Subsequently, membranes were extensively washed with 1x PBST, incubated with HyGLO chemiluminescent HRP antibody detection reagent (Denville), and imaged on an Amersham™ Imager 600 (GE Healthcare).

### Nonhomologous end joining (NHEJ) assay

Cell-free Trim-Away of the NHEJ factor PNKP in HSS was performed using 375 nM TRIM21^R-R-PS^ and 187.5 nM PNKP antibody (final concentrations) for 2 h at room temperature as described above. After this reaction, circular carrier DNA (pBlueScript) was added to all conditions and, where indicated, HSS was supplemented with recombinant *X. laevis* PNKP (a generous gift from Ben Stinson) to a final concentration of 50 nM. Samples for western blotting (see above) were withdrawn without further incubation. The ensemble NHEJ assay was essentially performed as described previously (Stinson et al., 2020). In short, joining reactions were initiated by the addition of radiolabeled linear DNA substrate (kindly provided by Ben Stinson) to 1 ng/µl final concentration and carried out at room temperature. At indicated times, samples were mixed with two volumes of clear agarose stop solution (80 mM Tris [pH 8.0], 8 mM EDTA, 0.13% phosphoric acid, 10% Ficoll, and 5% SDS) and digested with 1.25 mg/ml Proteinase K (Roche) for 60 min at 37°C. The DNA products were then separated by agarose gel electrophoresis on a native 0.7% agarose gel in 1x TBE. Gels were dried under vacuum, exposed to phosphor screens, and imaged on a Typhoon™ FLA 7000 phosphorimager (GE Healthcare).

### Replication-coupled DNA interstrand cross-link repair assay

Cell-free Trim-Away of TRAIP in both HSS and NPE was performed using 500 nM TRIM21^R-R-PS^ and each 125 nM of TRAIP-I and TRAIP-C antibodies (final concentrations) for 2 h at room temperature as described above. After this reaction, NPE of the indicated condition was supplemented with *X. laevis* TRAIP that was produced in the TnT® SP6 High-Yield Wheat Germ Protein Expression System (Promega) according to manufacturer’s instructions. Samples for western blotting (see above) were withdrawn from NPE without further incubation. Replication-coupled DNA interstrand cross-link (ICL) repair was essentially performed as described previously (Wu et al., 2019). In short, DNA damage-containing pICL (kindly provided by Alex Wu) was licensed at a final concentration of 7.5 ng/µl for 20 min at room temperature in HSS supplemented with trace amounts of [*α*-^32^P]dATP (Perkin Elmer). DNA replication was initiated by addition of two volumes of NPE mixture to one volume of licensing mixture. At indicated times, reactions were stopped with six volumes of replication stop solution (80 mM Tris [pH 8.0], 8 mM EDTA, 0.13% phosphoric acid, 10% Ficoll, 5% SDS, and 0.2% bromophenol blue) and treated with 3 mg/ml Proteinase K (Roche) for 1 h at 37°C. Replication intermediates were resolved on a native 0.9% agarose gel in 1x TBE. The gel was dried under vacuum, exposed to a phosphor screen, and imaged on a Typhoon™ FLA 7000 phosphorimager (GE Healthcare).

### Ubiquitin chain restriction (UbiCRest) analysis

Cell-free Trim-Away reactions of immobilized target proteins were performed as described above. After the reactions and high-salt washes, the beads were extensively washed with DUB reaction buffer (50 mM Tris [pH 7.5], 50 mM NaCl, and 5 mM DTT) supplemented with 0.02% Tween-20 and finally resuspended in DUB reaction buffer without Tween-20. The preparation of in vitro assembled polyubiquitin chains and ubiquitinated non-immobilized targets subjected to UbiCRest analysis is described below. UbiCRest analysis was essentially performed as described previously (Hospenthal et al., 2015). DUBs were diluted in DUB reaction buffer to 10x final concentrations containing 10 µM USP2 (Bio-Techne), 50 µM OTUB1*, 50 µM AMSH*, or both OTUB1* and AMSH* (50 µM each). Reactions were started by the addition of 10x concentrated DUB stocks to the samples and incubated for 1 h at 37°C on a rotating wheel. Reactions were stopped and subjected to SDS-PAGE and western blotting as described above.

### In vitro ubiquitination assays

Reconstituted ubiquitination reactions were prepared with 10x ubiquitination buffer (400 mM Tris [pH 7.5], 100 mM MgCl_2_, and 6 mM DTT) and contained final concentrations of 10 mM ATP, 50 µM ubiquitin, 100 nM E1 enzyme (Bio-Techne), 500 nM of each indicated E2 enzyme (UBE2W, UBE2N/UBE2V1, and/or UBE2D2; Bio-Techne), and 500 nM of the indicated TRIM21^R-R-PS^ or TRIM21^FL^ variant unless otherwise indicated. Reactions of antibody-dependent target ubiquitination additionally contained 100 nM recombinant substrate and, where indicated, 250 nM antibody. All reactions were incubated at 37°C for indicated times, stopped with 2x Laemmli sample buffer, and analyzed by SDS-PAGE and western blotting (see above). Reactions for UbiCRest analysis contained 250 nM TRIM21^FL^ in addition to the components described above, and were incubated for 20 min at 37°C. Reactions were stopped with an equal volume of dilution buffer (60 mM Tris [pH 7.5], 100 mM NaCl, and 10 mM DTT) supplemented with 25 mU/µl apyrase (NEB), and incubated for 10 min at 30°C. UbiCRest analysis was then performed as outlined above.

## REFERENCES

Beck, M., Schmidt, A., Malmstroem, J., Claassen, M., Ori, A., Szymborska, A., Herzog, F., Rinner, O., Ellenberg, J., and Aebersold, R. (2011). The quantitative proteome of a human cell line. Mol Syst Biol 7, 549–549.

Chen, X., Liu, M., Lou, H., Lu, Y., Zhou, M.-T., Ou, R., Xu, Y., and Tang, K.-F. (2019). Degradation of endogenous proteins and generation of a null-like phenotype in zebrafish using Trim-Away technology. Genome Biol 20, 19.

Clift, D., McEwan, W.A., Labzin, L.I., Konieczny, V., Mogessie, B., James, L.C., and Schuh, M. (2017). A Method for the Acute and Rapid Degradation of Endogenous Proteins. Cell 171, 1692–1706.e18.

Clift, D., So, C., McEwan, W.A., James, L.C., and Schuh, M. (2018). Acute and rapid degradation of endogenous proteins by Trim-Away. Nat Protoc 13, 2149–2175.

Dewar, J.M., Low, E., Mann, M., Räschle, M., and Walter, J.C. (2017). CRL2Lrr1 promotes unloading of the vertebrate replisome from chromatin during replication termination. Gene Dev 31, 275–290.

Duxin, J.P., Dewar, J.M., Yardimci, H., and Walter, J.C. (2014). Repair of a DNA-Protein Crosslink by Replication-Coupled Proteolysis. Cell 159, 346–357.

Fletcher, A.J., Mallery, D.L., Watkinson, R.E., Dickson, C.F., and James, L.C. (2015). Sequential ubiquitination and deubiquitination enzymes synchronize the dual sensor and effector functions of TRIM21. Proc National Acad Sci 112, 10014–10019.

Gillespie, P.J., Gambus, A., and Blow, J.J. (2012). Preparation and use of Xenopus egg extracts to study DNA replication and chromatin associated proteins. Methods San Diego Calif 57, 203–213.

Hardwick, L.J.A., and Philpott, A. (2015). An oncologist?s friend: How Xenopus contributes to cancer research. Dev Biol 408, 180–187.

Hauler, F., Mallery, D.L., McEwan, W.A., Bidgood, S.R., and James, L.C. (2012). AAA ATPase p97/VCP is essential for TRIM21-mediated virus neutralization. Proc National Acad Sci 109, 19733–19738.

Heubes, S., and Stemmann, O. (2007). The AAA-ATPase p97-Ufd1-Npl4 is required for ERAD but not for spindle disassembly in Xenopus egg extracts. J Cell Sci 120, 1325–1329.

Hoogenboom, W.S., Douwel, D.K., and Knipscheer, P. (2017). Xenopus egg extract: A powerful tool to study genome maintenance mechanisms. Dev Biol 428, 300–309.

Hospenthal, M.K., Mevissen, T.E.T., and Komander, D. (2015). Deubiquitinase-based analysis of ubiquitin chain architecture using Ubiquitin Chain Restriction (UbiCRest). Nat Protoc 10, 349–361.

Israel, S., Casser, E., Drexler, H.C.A., Fuellen, G., and Boiani, M. (2019). A framework for TRIM21-mediated protein depletion in early mouse embryos: recapitulation of Tead4 null phenotype over three days. BMC Genomics 20, 755.

James, L.C., Keeble, A.H., Khan, Z., Rhodes, D.A., and Trowsdale, J. (2007). Structural basis for PRYSPRY-mediated tripartite motif (TRIM) protein function. Proc National Acad Sci 104, 6200–6205.

Kiss, L., and James, L.C. (2021). The molecular mechanisms that drive intracellular neutralization by the antibody-receptor and RING E3 ligase TRIM21. Semin Cell Dev Biol 126, 99–107.

Kiss, L., Zeng, J., Dickson, C.F., Mallery, D.L., Yang, J.-C., McLaughlin, S.H., Boland, A., Neuhaus, D., and James, L.C. (2019). A tri-ionic anchor mechanism drives Ube2N-specific recruitment and K63-chain ubiquitination in TRIM ligases. Nat Commun 10, 4502.

Kiss, L., Clift, D., Renner, N., Neuhaus, D., and James, L.C. (2021). RING domains act as both substrate and enzyme in a catalytic arrangement to drive self-anchored ubiquitination. Nat Commun 12, 1220.

Larsen, N.B., Gao, A.O., Sparks, J.L., Gallina, I., Wu, R.A., Mann, M., Räschle, M., Walter, J.C., and Duxin, J.P. (2019). Replication-Coupled DNA-Protein Crosslink Repair by SPRTN and the Proteasome in Xenopus Egg Extracts. Mol Cell 73, 574–588.e7.

Mallery, D.L., McEwan, W.A., Bidgood, S.R., Towers, G.J., Johnson, C.M., and James, L.C. (2010). Antibodies mediate intracellular immunity through tripartite motif-containing 21 (TRIM21). Proc National Acad Sci 107, 19985–19990.

McEwan, W.A., Tam, J.C.H., Watkinson, R.E., Bidgood, S.R., Mallery, D.L., and James, L.C. (2013). Intracellular antibody-bound pathogens stimulate immune signaling via Fc-receptor TRIM21. Nat Immunol 14, 327–336.

Mevissen, T.E.T., and Komander, D. (2017). Mechanisms of Deubiquitinase Specificity and Regulation. Annu Rev Biochem 86, 1–33.

Mevissen, T.E.T., Hospenthal, M.K., Geurink, P.P., Elliott, P.R., Akutsu, M., Arnaudo, N., Ekkebus, R., Kulathu, Y., Wauer, T., Oualid, F.E., et al. (2013). OTU Deubiquitinases Reveal Mechanisms of Linkage Specificity and Enable Ubiquitin Chain Restriction Analysis. Cell 154, 169–184.

Michel, M.A., Elliott, P.R., Swatek, K.N., Simicek, M., Pruneda, J.N., Wagstaff, J.L., Freund, S.M.V., and Komander, D. (2015). Assembly and Specific Recognition of K29- and K33-Linked Polyubiquitin. Mol Cell 58, 95–109.

Räschle, M., Knipscheer, P., Knipsheer, P., Enoiu, M., Angelov, T., Sun, J., Griffith, J.D., Ellenberger, T.E., Schärer, O.D., and Walter, J.C. (2008). Mechanism of Replication-Coupled DNA Interstrand Crosslink Repair. Cell 134, 969–980.

Romano, F.B., Blok, N.B., and Rapoport, T.A. (2019). Peroxisome protein import recapitulated in Xenopus egg extracts. J Cell Biology 218, 2021–2034.

Scaglione, K.M., Basrur, V., Ashraf, N.S., Konen, J.R., Elenitoba-Johnson, K.S.J., Todi, S.V., and Paulson, H.L. (2013). The Ubiquitin-conjugating Enzyme (E2) Ube2w Ubiquitinates the N Terminus of Substrates*. J Biol Chem 288, 18784–18788.

Semlow, D.R., and Walter, J.C. (2021). Mechanisms of Vertebrate DNA Interstrand Cross-Link Repair. Annu Rev Biochem 90, 1–29.

Sparks, J., and Walter, J.C. (2019). Extracts for Analysis of DNA Replication in a Nucleus-Free System. Cold Spring Harb Protoc 2019, pdb.prot097154.

Stinson, B.M., and Loparo, J.J. (2021). Repair of DNA Double-Strand Breaks by the Nonhomologous End Joining Pathway. Annu Rev Biochem 90, 1–28.

Stinson, B.M., Moreno, A.T., Walter, J.C., and Loparo, J.J. (2020). A Mechanism to Minimize Errors during Non-homologous End Joining. Mol Cell 77, 1080–1091.e8.

Swatek, K.N., Usher, J.L., Kueck, A.F., Gladkova, C., Mevissen, T.E.T., Pruneda, J.N., Skern, T., and Komander, D. (2019). Insights into ubiquitin chain architecture using Ub-clipping. Nature 572, 533–537.

Tatham, M.H., Plechanovová, A., Jaffray, E.G., Salmen, H., and Hay, R.T. (2013). Ube2W conjugates ubiquitin to α-amino groups of protein N-termini. Biochem J 453, 137–145.

Twomey, E.C., Ji, Z., Wales, T.E., Bodnar, N.O., Ficarro, S.B., Marto, J.A., Engen, J.R., and Rapoport, T.A. (2019). Substrate processing by the Cdc48 ATPase complex is initiated by ubiquitin unfolding. Science 365, eaax1033.

Vittal, V., Shi, L., Wenzel, D.M., Scaglione, K.M., Duncan, E.D., Basrur, V., Elenitoba-Johnson, K.S.J., Baker, D., Paulson, H.L., Brzovic, P.S., et al. (2015). Intrinsic disorder drives N-terminal ubiquitination by Ube2w. Nat Chem Biol 11, 83–89.

Wauer, T., Swatek, K.N., Wagstaff, J.L., Gladkova, C., Pruneda, J.N., Michel, M.A., Gersch, M., Johnson, C.M., Freund, S.M., and Komander, D. (2015). Ubiquitin Ser65 phosphorylation affects ubiquitin structure, chain assembly and hydrolysis. Embo J 34, 307–325.

Weir, E., McLinden, G., Alfandari, D., and Cousin, H. (2021). Trim-Away mediated knock down uncovers a new function for Lbh during gastrulation of Xenopus laevis. Dev Biol 470, 74–83.

Wu, R.A., Semlow, D.R., Kamimae-Lanning, A.N., Kochenova, O.V., Chistol, G., Hodskinson, M.R., Amunugama, R., Sparks, J.L., Wang, M., Deng, L., et al. (2019). TRAIP is a master regulator of DNA interstrand cross-link repair. Nature 567, 267–272.

Wühr, M., Freeman, R.M., Presler, M., Horb, M.E., Peshkin, L., Gygi, S.P., and Kirschner, M.W. (2014). Deep Proteomics of the Xenopus laevis Egg using an mRNA-Derived Reference Database. Curr Biol 24, 1467–1475.

Zeng, J., Santos, A.F., Mukadam, A.S., Osswald, M., Jacques, D.A., Dickson, C.F., McLaughlin, S.H., Johnson, C.M., Kiss, L., Luptak, J., et al. (2021). Target-induced clustering activates Trim-Away of pathogens and proteins. Nat Struct Mol Biol 28, 278–289.

