## Supplemental Figures for "Cell-free Trim-Away reveals the mechanism of antibody-mediated protein degradation by TRIM21"

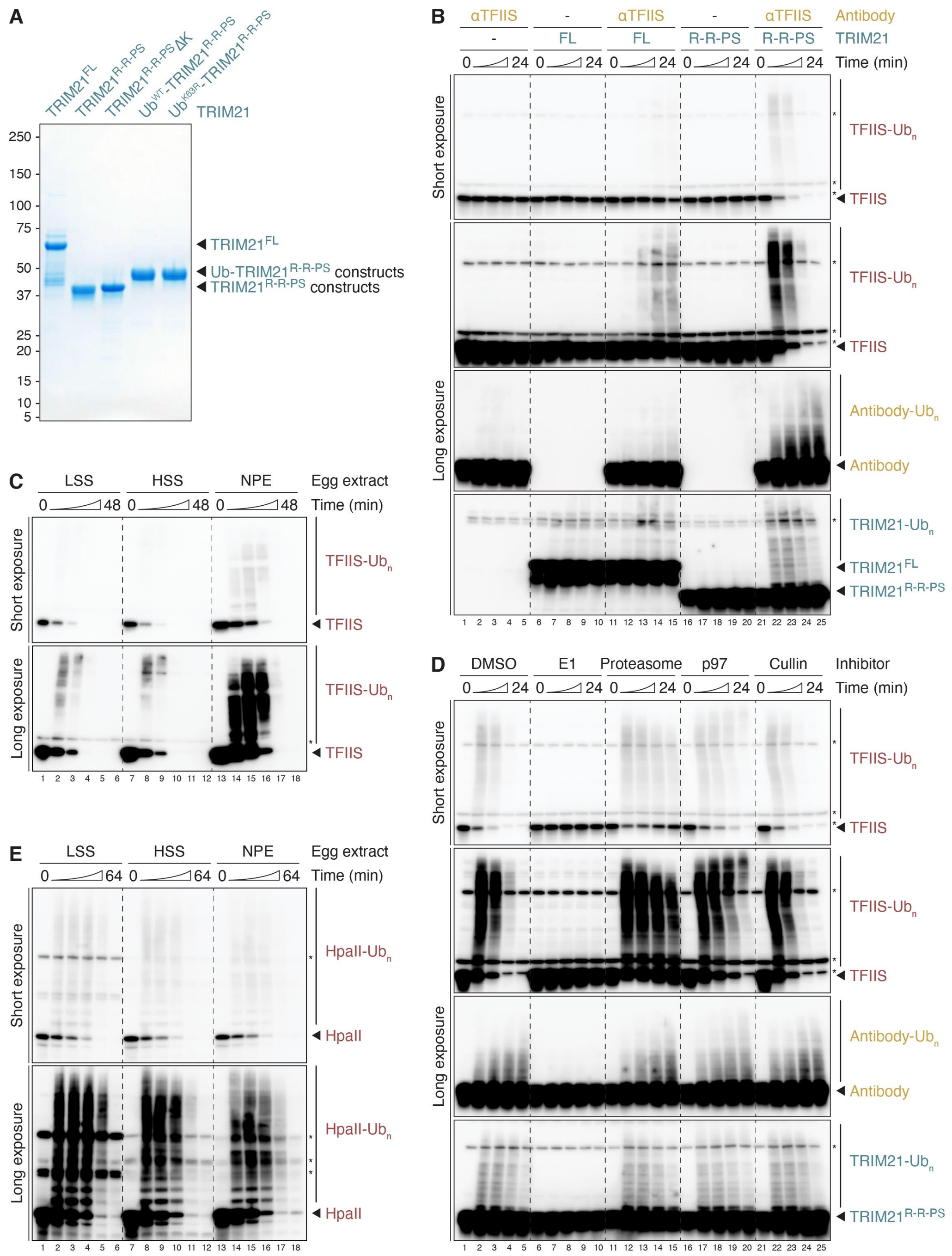


**Figure S1. Cell-free Trim-Away in *Xenopus* egg extracts. (A)** Coomassie staining of purified recombinant human TRIM21 variants used in this study. **(B)** Trim-Away of endogenous TFIIS in HSS (high-speed supernatant) comparing $TRIM21^{FL}$ and $TRIM21^{R\text{-}R\text{-}PS}$. **(C)** Endogenous TFIIS was targeted by $TRIM21^{R\text{-}R\text{-}PS}$ in three types of egg extract: LSS (low-speed supernatant), HSS (high-speed supernatant), and NPE (nucleoplasmic extract). Note that TFIIS is more abundant in NPE than in the other extracts. **(D)** Trim-Away of TFIIS in HSS in the presence of inhibitors targeting E1 enzyme (MLN7243), the proteasome (MG-262), p97 (NMS-873), or the NEDD8 activating enzyme to inhibit Cullin-RING E3 ligases (MLN4924). **(E)** Egg extracts were supplemented with 250 nM recombinant HpaII to compare Trim-Away efficiencies using $TRIM21^{R\text{-}R\text{-}PS}$ in LSS, HSS, and NPE at a constant target protein concentration.

Samples in panels B-E were stopped at indicated times and directly analyzed by SDS-PAGE and western blotting. In contrast, samples of corresponding experiments shown in Figures 1C-1F were treated with E1 and proteasome inhibitors as well as the non-specific deubiquitinase USP2 at indicated times for 30 min prior to SDS-PAGE and western blot analysis. This treatment collapsed all ubiquitin chains and preserved the remaining Trim-Away components for better assessment of degradation efficiencies. Unmodified proteins of interest are highlighted by an arrow. We assume that any bands already present at the start of the reaction are non-specific (asterisk).

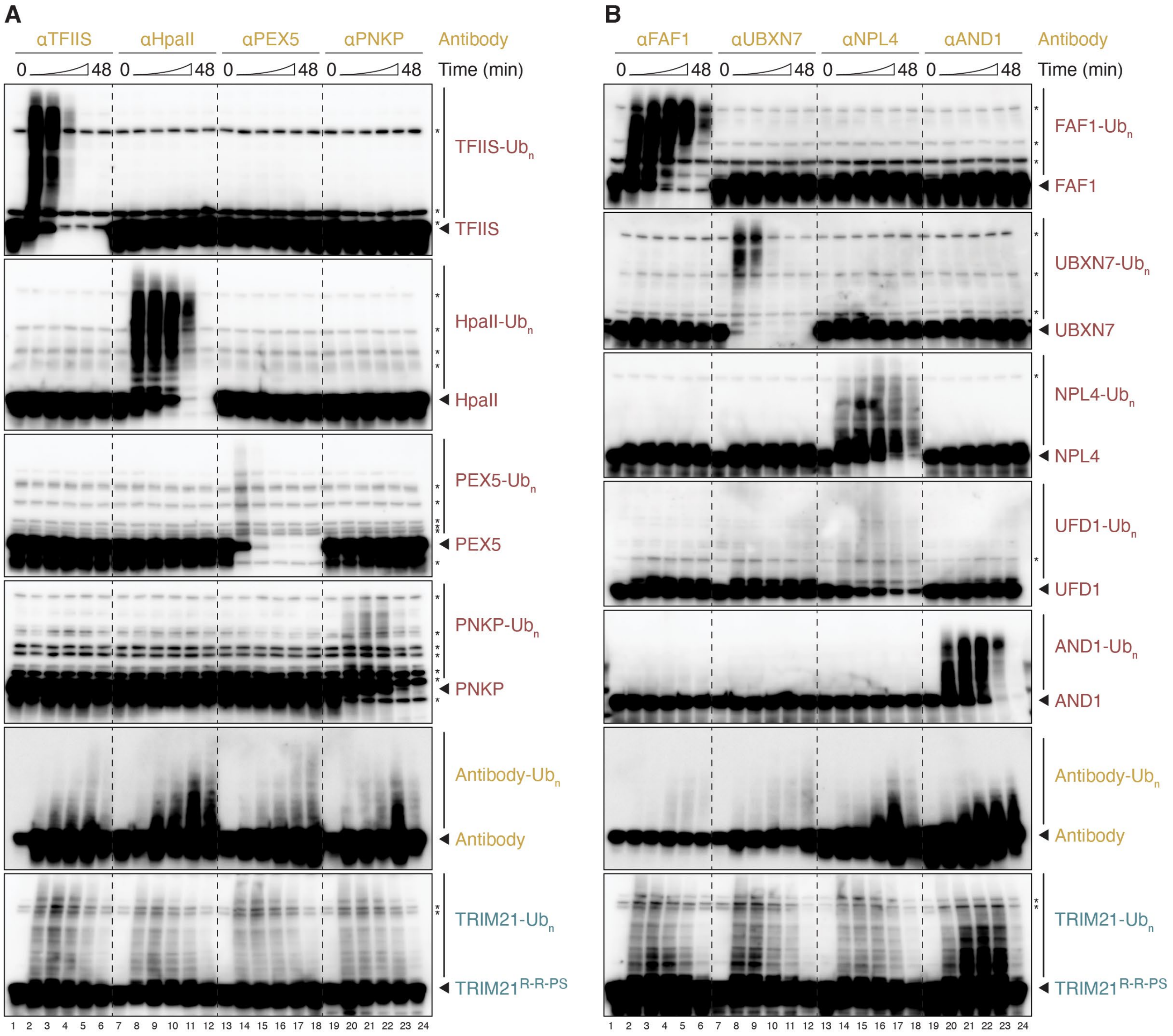


**Figure S2. All three components are directly ubiquitinated during cell-free Trim-Away. (A, B)** The fate of target proteins during cell-free Trim-Away prior to proteasomal degradation was assessed in HSS. Comparison of the same panel of endogenous and exogenous substrates targeted in Figure 1G. Samples were taken over a time course (0, 3, 6, 12, 24, and 48 min) prior to SDS-PAGE and western blot analysis. Note that UFD1 is a constitutive binding partner of the targeted protein NPL4. Shorter exposures of the same western blots, except for antibody and TRIM21, are shown in Figure 3.

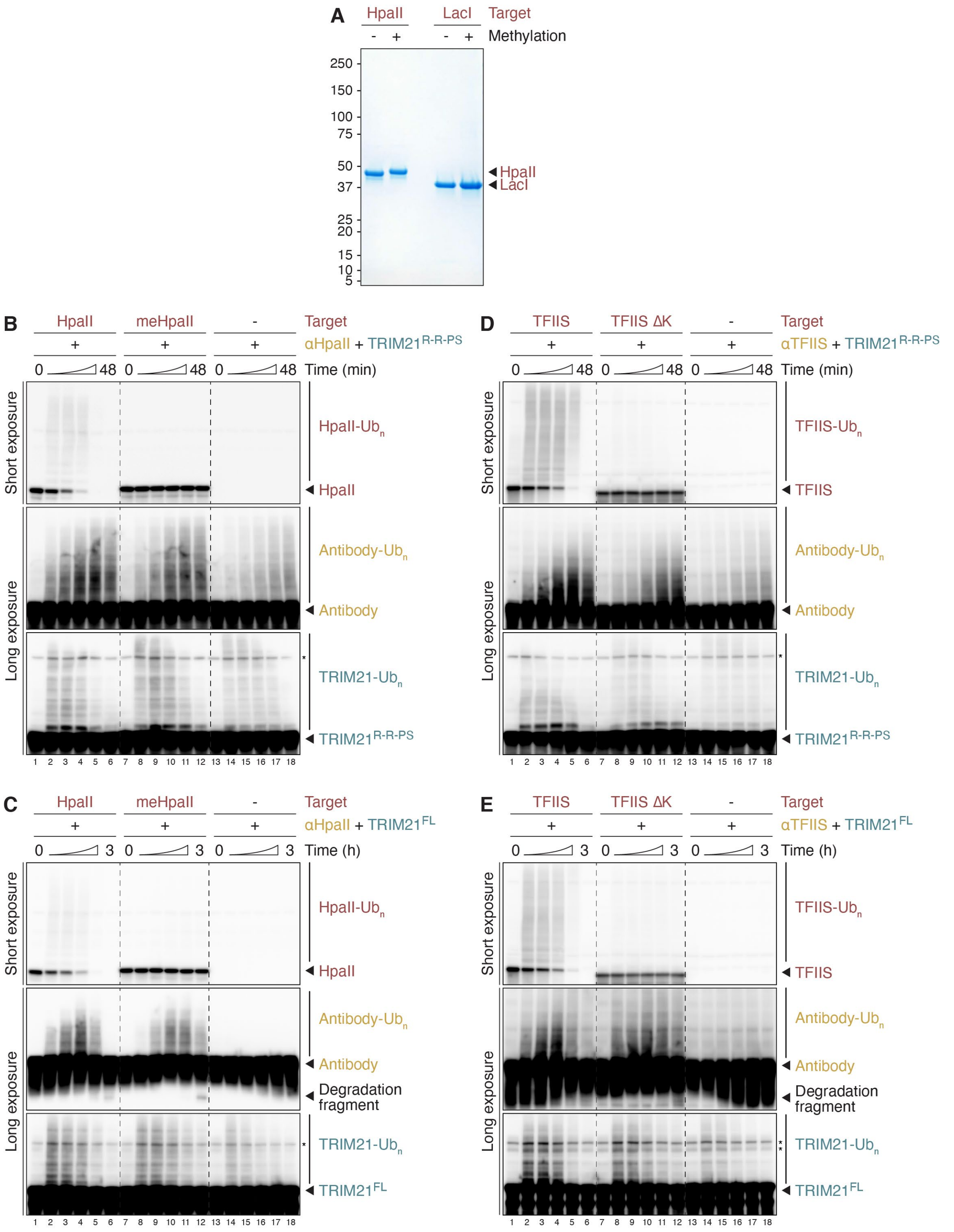


**Figure S3. Direct polyubiquitination of targets is essential for their destruction. (A)** Coomassie staining of recombinant HpaII and LacI proteins before and after reductive methylation. Note that successful methylation results in a slight gel shift. **(B, C)** Trim-Away of recombinant unmethylated or methylated HpaII (final concentration of 250 nM) by TRIM21$^{R\text{-}R\text{-}PS}$ (B) or TRIM21$^{FL}$ (C) in HSS. **(D, E)** Trim-Away of recombinant TFIIS variants (final concentration of 200 nM) by TRIM21$^{R\text{-}R\text{-}PS}$ (D) or TRIM21$^{FL}$ (E) in HSS. Endogenous TFIIS was immunodepleted from egg extract prior to the addition of recombinant *X. laevis* TFIIS. Trim-Away with TRIM21$^{FL}$ required an extended time course (0, 11, 22, 45, 90, and 180 min) for efficient HpaII (C) and TFIIS (E) degradation.

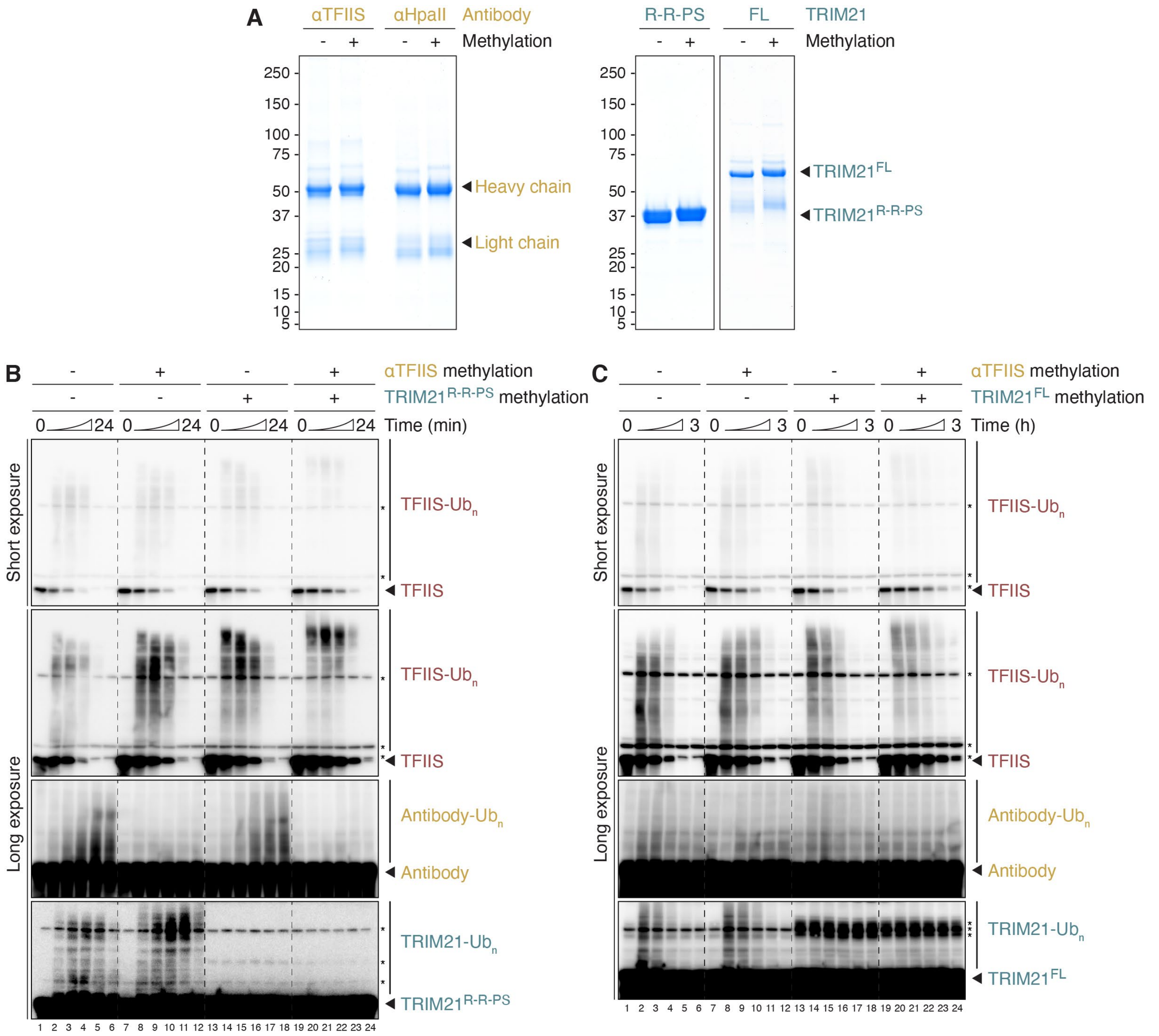


**Figure S4. Direct target polyubiquitination is sufficient for Trim-Away. (A)** Coomassie staining of TFIIS and HpaII antibodies (left panel) and TRIM21 variants (right panel) before and after reductive methylation. Note that successful methylation results in a slight gel shift. **(B, C)** Cell-free Trim-Away of endogenous TFIIS by TFIIS antibody and TRIM21$^{R-R-PS}$ (B) or TRIM21$^{FL}$ (C). The methylation state of TFIIS antibody and TRIM21 is indicated. Trim-Away with TRIM21$^{FL}$ required an extended time course (0, 11, 22, 45, 90, and 180 min) for efficient TFIIS degradation. Note that methylation of TRIM21$^{FL}$ resulted in an even stronger non-specific band (asterisks), which is already present at the beginning of the Trim-Away reaction.

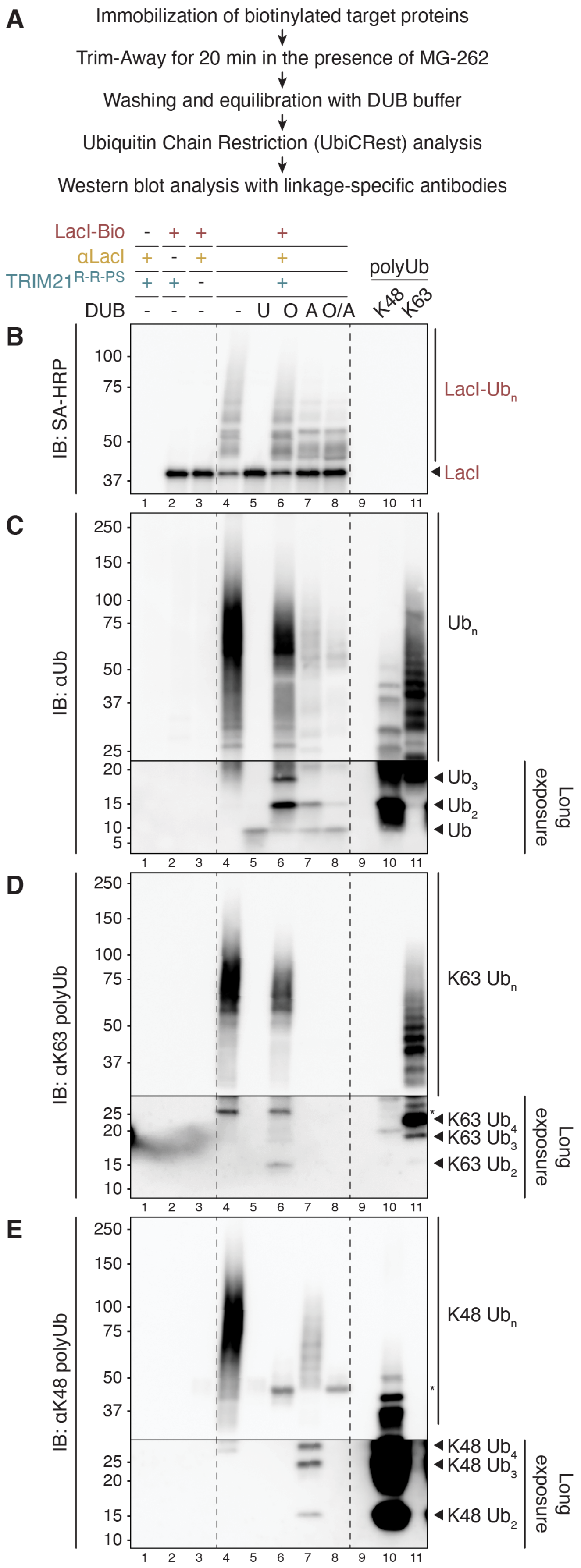


**Figure S5. UbiCRest analysis reveals heterotypic K48/K63 ubiquitin chains on Trim-Away targets.** **(A)** Experimental strategy for the characterization of substrate ubiquitin chain architecture by Ubiquitin Chain Restriction (UbiCRest) analysis and linkage-specific ubiquitin antibodies. **(B-E)** Analysis of antibody- and TRIM21-mediated polyubiquitination of immobilized biotinylated LacI. After Trim-Away in HSS containing proteasome inhibitor (MG-262), the streptavidin-coupled magnetic beads were washed and treated with the following deubiquitinases (DUBs): non-specific USP2, K48 linkage-specific OTUB1*, K63-specific AMSH*, or both OTUB1* and AMSH*. Treated samples were analyzed by SDS-PAGE and western blotting, visualizing biotinylated LacI (B), total ubiquitin (C), K63-linked ubiquitin chains (D), or K48-linked ubiquitin chains (E). In lanes 10 and 11, recombinant K48- and K63-linked ubiquitin chains were loaded to confirm the specificities of ubiquitin antibodies and for comparing released polyubiquitin species after DUB treatment. The low-molecular weight sections in panels C-E were exposed longer than the high-molecular weight regions to support the identification of free ubiquitin species. Asterisks correspond to cross-reacting bands.

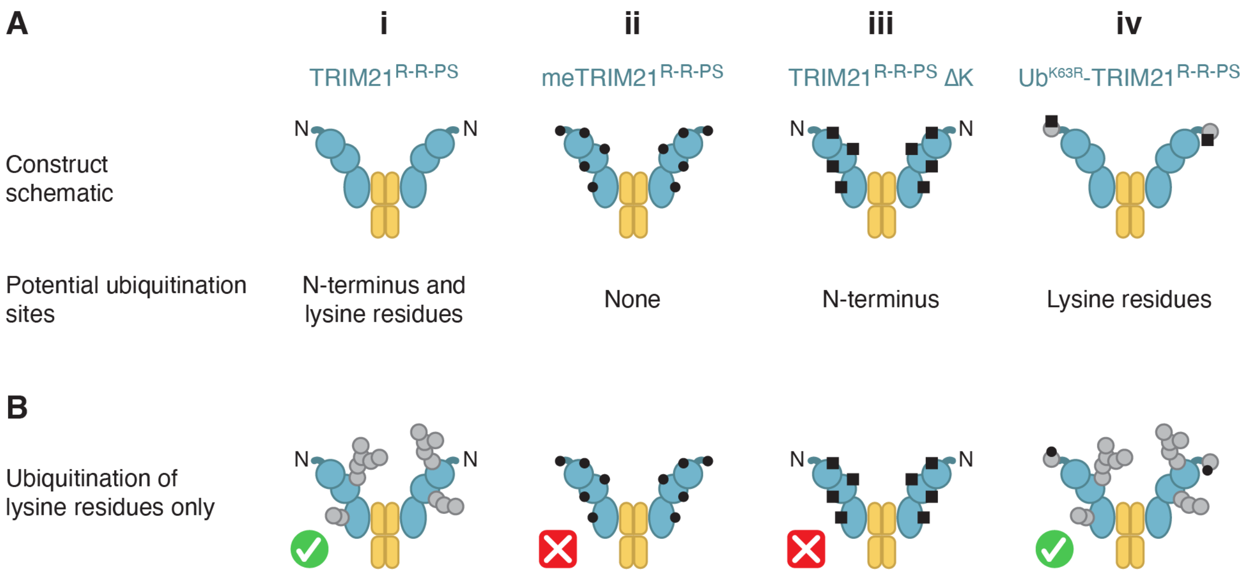


**Figure S6. TRIM21 is ubiquitinated on lysine residues during cell-free Trim-Away. (A)** Construct schematic and potential ubiquitination sites of i) TRIM21$^{R\text{-}R\text{-}PS}$ wild-type, ii) methylated TRIM21$^{R\text{-}R\text{-}PS}$, iii) lysine-less TRIM21$^{R\text{-}R\text{-}PS}$ (ΔK), and iv) a ubiquitin (K63R)-TRIM21$^{R\text{-}R\text{-}PS}$ fusion protein. **(B)** Schematic models of the TRIM21$^{R\text{-}R\text{-}PS}$ constructs in (A) after ubiquitination on lysines. These schematics are consistent with the TRIM21 ubiquitination pattern observed in cell-free Trim-Away (Figure 7B, bottom panel), suggesting that TRIM21 is ubiquitinated on lysines rather than its N-terminus. Available N-termini (N), methylated sites (black circle), and lysine to arginine mutations (black square) are highlighted.

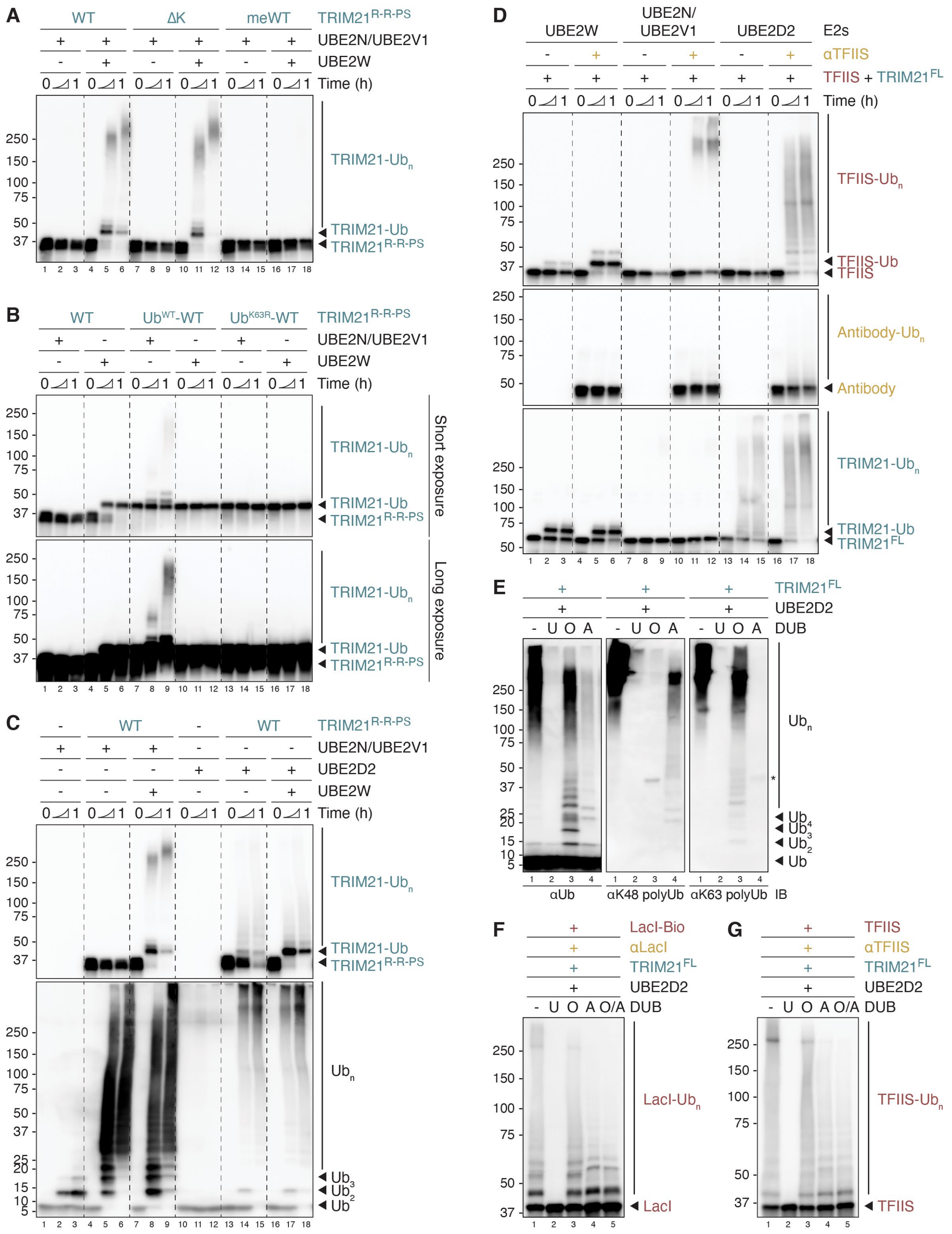


**Figure S7. In vitro validation of TRIM21 variants and reconstitution of antibody-dependent target ubiquitination by TRIM21. (A)** In vitro autoubiquitination assay comparing wild-type (WT), lysine-less (ΔK), and methylated TRIM21$^{R\text{-}R\text{-}PS}$ in reactions containing UBE2N/UBE2V1 without or with UBE2W. As expected, N-terminal TRIM21 ubiquitination was observed for WT and ΔK constructs, but not methylated TRIM21. **(B)** TRIM21 autoubiquitination assay comparing TRIM21$^{R\text{-}R\text{-}PS}$ WT and constructs featuring ubiquitin (WT or K63R) fused to the TRIM21 N-terminus. In the presence of UBE2N/UBE2V1, only Ub$^{WT}$-TRIM21$^{R\text{-}R\text{-}PS}$ was modified, but not TRIM21$^{R\text{-}R\text{-}PS}$ WT, which requires prior N-terminal monoubiquitination by UBE2W. The fact that Ub$^{K63R}$-TRIM21$^{R-R-PS}$ was not modified by UBE2N/UBE2V1 confirms that the only ubiquitination site was absent. Importantly, both N-terminal ubiquitin fusion proteins were not monoubiquitinated in the presence of UBE2W, as expected (Scaglione et al., 2013; Tatham et al., 2013; Vittal et al., 2015). Ub$^{K63R}$-TRIM21$^{R-R-PS}$ is therefore a variant with a blocked N-terminus (i.e. UBE2W and UBE2N/UBE2V1 cannot modify this construct), while all remaining lysines are available for ubiquitination. **(C)** In vitro ubiquitination reactions with TRIM21$^{R-R-PS}$ comparing UBE2N/UBE2V1 and UBE2D2 in the absence or presence of UBE2W. TRIM21 autoubiquitination (top panel) and free ubiquitin chain formation (bottom panel) are shown. UBE2N/UBE2V1 inefficiently assembled unattached K63-linked chains in the absence of an E3 ligase. TRIM21 strongly enhanced this activity independently of UBE2W. TRIM21 autoubiquitination by UBE2N/UBE2V1 required UBE2W-mediated N-terminal monoubiquitination, as seen in (A) and (B). While UBE2D2 alone did not assemble free ubiquitin chains, TRIM21 addition promoted efficient ubiquitin chain formation. Interestingly, TRIM21$^{R-R-PS}$ ubiquitinated itself to some extent when UBE2D2 was present. Addition of UBE2W did not enhance this modification, but instead all unmodified TRIM21 shifted up to a monoubiquitinated species. This data indicates that UBE2D2-mediated TRIM21 ubiquitination occurs on lysines, and is therefore independent of UBE2W activity. **(D)** Reconstitution of antibody-dependent target ubiquitination by TRIM21. Recombinant TFIIS was incubated with ATP, ubiquitin, E1, indicated E2 enzymes, and TRIM21$^{FL}$ in the absence or presence of TFIIS antibody. **(E)** UbiCRest analysis of polyubiquitin chains assembled with TRIM21$^{FL}$ and UBE2D2. In vitro reactions were treated with the following deubiquitinases (DUBs): non-specific USP2, K48 linkage-specific OTUB1*, or K63-specific AMSH*. Treated samples were analyzed by SDS-PAGE and western blotting, visualizing total ubiquitin (left panel), K48-linked ubiquitin chains (middle panel), or K63-linked ubiquitin chains (right panel). **(F-G)** UbiCRest analysis of LacI (F) or TFIIS (G) after antibody-dependent in vitro ubiquitination by TRIM21$^{FL}$ and UBE2D2. Both target proteins were modified with K48- and K63-linked ubiquitin chains, closely resembling the polyubiquitination pattern of substrates during cell-free Trim-Away (Figures 6A and S5B).
